# Chronic NLRP3 inflammasome activation drives neutrophil brain entry and interactions with microglia

**DOI:** 10.64898/2026.04.22.720282

**Authors:** Lukas L. Skuja, Sophia M. Guldberg, David Joy, Jason C. Dugas, Neal S. Gould, Roni Chau, David Tatarakis, Isabel Becerra, Cynthia Chau, Connie Ha, David Huynh, Hoang N. Nguyen, Lily Sarrafha, Elizabeth W. Sun, Shan V. Andrews, Thomas Sandmann, Jung H. Suh, Robert G. Thorne, Pamela J. Lein, Kathryn M. Monroe, Gilbert Di Paolo

## Abstract

NOD-like receptor family pyrin domain-containing 3 (NLRP3) is a cytosolic regulator of an inflammasome-mediated innate immune response. In the central nervous system (CNS), NLRP3 inflammasome activation has been implicated in multiple neurodegenerative diseases, yet the mechanisms by which it contributes to disease remain unclear. Here, we investigated the CNS effects of chronic NLRP3 activation using a humanized *NLRP3* gain-of-function mouse model (*hNLRP3^D305N^*). Bulk brain analyses confirmed constitutive inflammasome activation, widespread cytokine induction, and the increased presence of blood-associated proteins suggestive of dysfunction at CNS border sites and the blood-brain barrier (BBB). Furthermore, cerebrospinal fluid (CSF) neurofilament light chain levels were elevated, indicating neuronal damage. Single-cell RNA-sequencing of CD45^+^ immune cells in the brain demonstrated that microglia adopt distinct reactive states and that peripheral immune cells infiltrate the CNS, with neutrophils emerging as the predominant infiltrating immune cell type. This finding was confirmed by untargeted bulk brain and CSF proteomics that also suggest neutrophil reactivity. Immunohistochemistry further revealed regional neutrophil entry into the brain parenchyma, concurrent with reactive microglia and engulfment of neutrophils, suggesting functional microglia-neutrophil interactions. Collectively, these findings establish a direct pathogenic role for the NLRP3 inflammasome in the CNS independent of other neurodegeneration-related disease pathologies.

## Introduction

The NOD-like receptor family pyrin domain-containing 3 (NLRP3) inflammasome is a fundamental innate immune multiprotein complex that can be deleterious when chronically activated, as observed in the central nervous system (CNS) across diverse neurodegenerative diseases^1–3^. NLRP3 has emerged as a promising therapeutic target, with pharmacologic inhibitors currently being investigated in the clinic for both systemic and CNS indications^4,5^. NLRP3 is primarily expressed by myeloid cells, such as brain-resident microglia, and canonically requires both a priming and activation step^6,7^. NLRP3 priming is typically triggered by toll-like receptor ligands and cytokine signaling via NF-ΚB, which induces the transcription of NLRP3 and pro-IL-1β, as well as mediates post-translational modifications to NLRP3 to promote its activation^8,9^. NLRP3 activation can occur in response to a variety of stimuli including microbial toxins, lysosomal permeabilization, mitochondrial dysfunction, damage-associated molecular patterns (DAMPs), along with pathological protein aggregates that are characteristic of neurodegenerative diseases, such as amyloid-β, tau, and α-synuclein^10,11^. Upon activation, NLRP3 undergoes self-oligomerization and recruits apoptosis-associated speck-like protein containing a caspase activation and recruitment domain (CARD) (ASC), which subsequently recruits pro-caspase-1 and induces its self-cleavage to yield active caspase-1^12^. Caspase-1 then cleaves pro-IL-1β and pro-IL-18 into the mature forms of these proinflammatory cytokines^13^. Active caspase-1 also cleaves gasdermin D (GSDMD), releasing its N-terminal domain that is transferred to the plasma membrane where it forms pores through which IL-1β and IL-18 can be secreted^14^. GSDMD pores can also lead to a form of cell death known as pyroptosis^15^. The contents of the pyroptotic cell are released into the extracellular space, including DAMPs that can induce NLRP3 pathway activation in other cells^16^. In the context of sterile inflammation, NLRP3 activation propagates inflammatory responses while eliminating activated immune cells via pyroptosis^2,16,17^.

Growing evidence implicates the NLRP3 inflammasome in neurodegeneration^1,3,18^. Studies in humans have demonstrated that IL-1β levels are elevated in the CNS of patients with various neurodegenerative diseases, along with other manifestations of NLRP3 inflammasome pathway activation such as ASC oligomerization, cleaved caspase-1 and GSDMD, as well as the release of mature IL-18^19–23^. Cellular and animal models of neurodegenerative diseases, such as Alzheimer’s disease^24–27^, Parkinson’s disease^22,28–30^, amyotrophic lateral sclerosis^31–33^, and multiple sclerosis^34–37^, have also implicated NLRP3, with NLRP3 ablation or pharmacologic inhibition showing neuroprotection in these models. However, despite evidence linking the NLRP3 inflammasome to neurodegenerative diseases, the consequences of direct NLRP3 activation in the CNS in the absence of additional stimuli, such as neuropathology, and whether this alone is sufficient to induce neurodegeneration, remain largely unknown.

Rare gain-of-function (GoF) mutations found in human *NLRP3* result in a group of autoinflammatory diseases known as cryopyrin-associated periodic syndrome (CAPS). These diseases share symptoms such as rash, episodic unprovoked fever, fatigue, and abdominal pain, with the most severe cases resulting in mental impairments and organ damage^38^. Although *NLRP3* GoF mouse models have been developed to study CAPS and patients often present neurological impairments such as fatigue, headache, and hearing loss^39^, the CNS effects of *NLRP3* GoF mutations have not been thoroughly investigated. Previous studies have described increased blood-brain barrier (BBB) permeability and neutrophil CNS infiltration in a neutrophil-specific *Nlrp3* GoF mouse model^40^. Additionally, humanized *NLRP3* mice with the D305N GoF mutation (*hNLRP3^D305N^*) display constitutive systemic inflammation, meningitis, and increased expression of select inflammation-associated genes in bulk brain homogenate^41–44^. Importantly, meningeal peripheral immune cell profiling has largely relied on cell surface marker analysis by flow cytometry that does not elucidate whether these cells enter the brain parenchyma to interact with CNS-resident myeloid cells, such as microglia, and overall CNS characterization remains limited.

Here, we utilized *hNLRP3^D305N^* mice to investigate the impact of chronic NLRP3 activation on the CNS. We identified features of NLRP3 pathway activity, neuroinflammation, BBB permeability, and neuronal damage in these mice without exogenous stimuli. Single-cell RNA sequencing (scRNA-seq) of microglia demonstrated broad transcriptional shifts towards more reactive states and the emergence of distinct subpopulations that were also defined by expression of proinflammatory genes. Transcriptomic analysis of all immune cell populations found in the brains of these mice revealed a robust accumulation of peripheral immune cell types, most strikingly neutrophils, that were almost absent in wild-type (WT) mice. This finding was confirmed by untargeted bulk brain and cerebrospinal fluid (CSF) proteomics that also suggest neutrophil reactivity. Immunohistochemical (IHC) staining of brain tissue confirmed immune cell infiltration in the leptomeninges and demonstrated regional neutrophil entry into the brain parenchyma, including in the septal nucleus, fimbria, and areas proximal to the leptomeninges. Super-resolution microscopy further revealed apparent interactions between neutrophils and microglia in these regions, as well as changes in microglial morphology and protein expression that are consistent with activated states. Together, these data indicate that the *hNLRP3^D305N^*GoF mutation drives microglial activation, neutrophil recruitment, and immune cell crosstalk that likely exacerbates neuroinflammation in the brain. Collectively, these findings demonstrate that chronic NLRP3 activation is sufficient to drive sterile neuroinflammation, BBB permeability, and neuronal damage, thereby establishing a direct pathogenic role for NLRP3 in the absence of additional disease-associated pathologies.

## Results

### The NLRP3 inflammasome is constitutively active in the brain and peripherally in *hNLRP3^D305N^* mice

Previous characterization of the *hNLRP3^D305N^* and *hNLRP3^WT^* mouse models demonstrated that *hNLRP3^D305N^*mice have features of peripheral inflammation under basal conditions, notably including elevated IL-1β and IL-1RA, the endogenous competitive antagonist of IL-1β that is often upregulated in tandem^41,45^. However, existing investigations into constitutive NLRP3 pathway activation in the CNS have relied on indirect markers of neuroinflammation and an IL-1β western blot that lacked sufficient power for statistical analysis^43^. We homogenized bulk brain samples from *hNLRP3^D305N^* mice, as well as their *mNlrp3^WT^* littermates and a *hNLRP3^WT^* line to control for the humanization of *NLRP3* and potential mouse strain differences. All mice characterized in this study were homozygous for *mNlrp3^WT^*, *hNLRP3^WT^*, or *hNLRP3^D305N^*. IL-1β and IL-1RA levels were significantly elevated in *hNLRP3^D305N^* brains compared to controls (Fig. 1a). The two control lines showed relatively similar IL-1β and IL-1RA levels, suggesting that humanization of *NLRP3* does not substantially alter basal NLRP3 pathway activity. Additional comparisons between *hNLRP3^D305N^* and *hNLRP3^WT^* mouse brains revealed elevated ASC by western blot in the GoF mutation carriers, further demonstrating constitutive NLRP3 inflammasome activation (Supplementary Fig. 1a-b). IL-18 levels were only modestly higher in *hNLRP3^D305N^*mouse brains compared to *hNLRP3^WT^* (Supplementary Fig. 1c), although plasma IL-18 levels, as well as plasma IL-1β and IL-1RA levels, were elevated in *hNLRP3^D305N^* as expected (Supplementary Fig. 1c-d). Altogether, these results demonstrate constitutive NLRP3 pathway activation in the brains and periphery of *hNLRP3^D305N^*mice.

**Figure 1.**
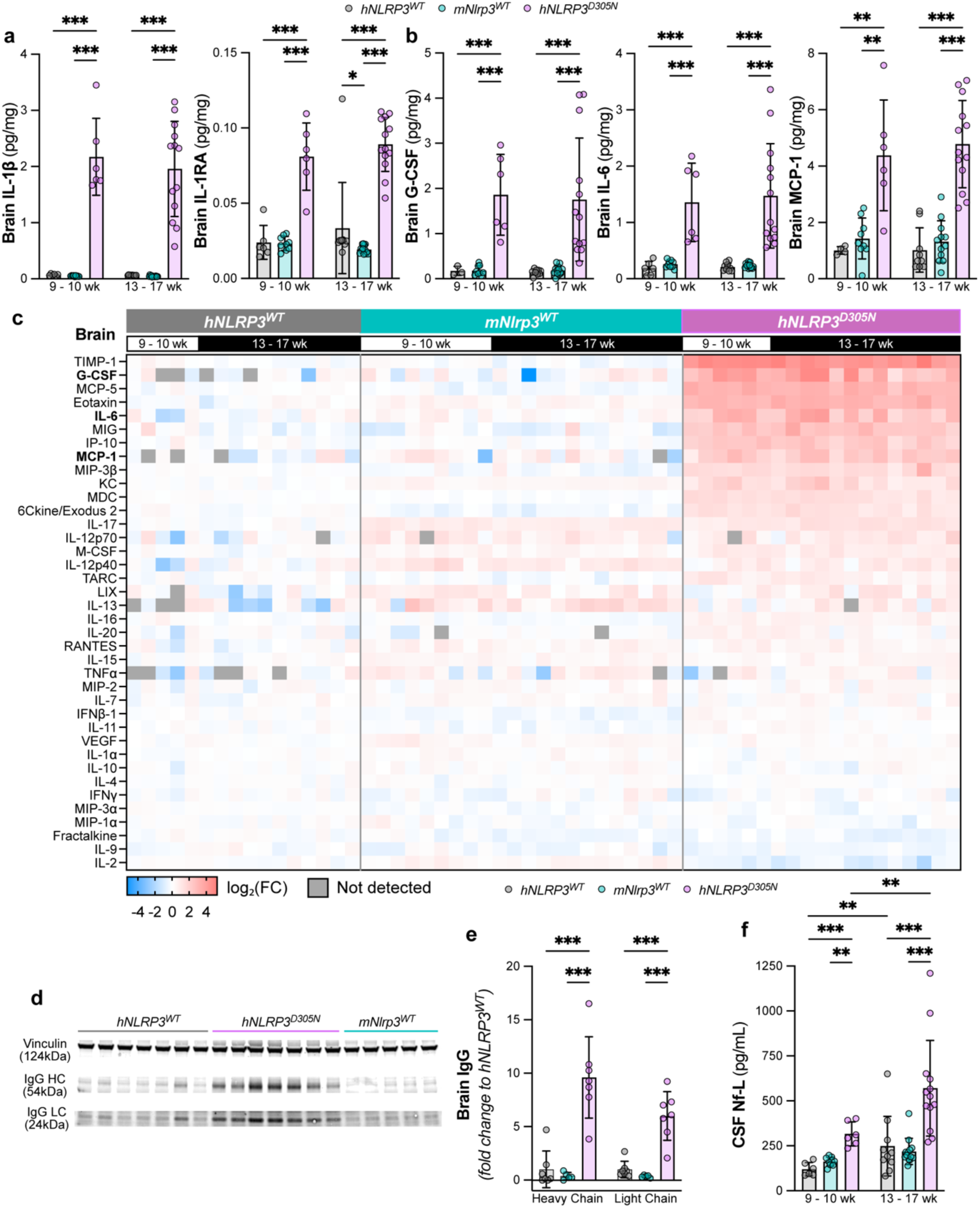
*hNLRP3^D305N^*mice show constitutive inflammasome activity in the brain, broad neuroinflammatory signaling, and signs of BBB permeability and neuronal damage (a) IL-1β MSD and IL-1RA ELISA quantification of bulk brain samples (n = 6-13/group). **(b-c)** Bulk brain samples were assayed in a cytokine panel. Select cytokines are shown in column graphs (n = 3-13/group) and the full panel is shown as heatmap (columns represent individual mice). Heatmap values are normalized to *hNLRP3^WT^*. **(d-e)** Representative images and quantification of western blot for IgG heavy chain (HC) and IgG light chain (LC) in bulk brain samples. Vinculin is used as loading control. Normal two-way ANOVA with Tukey multiple testing correction for comparisons across genotype (n = 5-7/group). **(f)** Nf-L Simoa assay quantification in CSF samples (n = 6-13/group). All panels include *hNLRP3^WT^*, *mNlrp3^WT^*, and *hNLRP3^D305N^* mice. Unless otherwise noted, lognormal two-way ANOVA with Tukey multiple testing correction for comparisons across genotype and age group. *p_adj_ < 0.05, **p_adj_ < 0.01, ***p_adj_ < 0.001. Data are shown as mean ± SD. Full source data and statistic values in Source Data.

### Constitutive NLRP3 activation induces broad neuroinflammatory signaling in *hNLRP3^D305N^* mice

With constitutive activation of NLRP3 in the brain, we hypothesized that broader neuroinflammation may occur in *hNLRP3^D305N^* mice. A previous study showed increased expression of a select number of inflammation-associated genes examined by RT-qPCR^42^. We therefore explored changes in a neuroinflammatory protein panel and identified broad proinflammatory cytokine and chemokine production in *hNLRP3^D305N^*mouse brains, including G-CSF, IL-6, and MCP-1 (Fig. 1b-c).

### Blood-associated proteins present in *hNLRP3^D305N^*mouse brains suggest increased blood-brain barrier permeability

NLRP3 activation has been shown to affect BBB permeability^40,46^, along with proinflammatory cytokines elevated in *hNLRP3^D305N^* mouse brains^47^. To evaluate BBB permeability in the *hNLRP3^D305N^* mice, we assessed endogenous immunoglobulin G (IgG) and serum albumin levels by western blot, finding both peripherally restricted proteins were elevated in *hNLRP3^D305N^*mouse brains at baseline (Fig. 1d-e; Supplementary Fig. 2a-b). We confirmed IgG levels were higher in *hNLRP3^D305N^* mouse brains and plasma compared to *hNLRP3^WT^* by MSD, consistent with systemic inflammation (Supplementary Fig. 2c). Notably, the increased brain:plasma ratio of IgG in *hNLRP3^D305N^* mice, along with a general increase in IHC signal across both cortical and subcortical brain regions (Supplementary Fig. 2d-e), suggests BBB permeability to IgG was likely increased in addition to global elevations of endogenous IgG. Brain IHC for albumin similarly revealed broad parenchymal increases in *hNLRP3^D305N^* mice compared to the WT controls (Supplementary Fig. 2d-e). These results suggest increased BBB permeability occurs in *hNLRP3^D305N^* mice.

### Neurofilament light chain is elevated in cerebrospinal fluid of *hNLRP3^D305N^*mice indicative of neuronal damage

While NLRP3 has been implicated in neurodegeneration, it remains unclear whether its activation alone is sufficient to drive neuronal damage. We therefore investigated neurofilament light chain (Nf-L) levels, a biomarker for neurodegeneration due to its release from neurons upon death or axonal damage^48,49^, in the CSF of *hNLRP3^WT^*, *mNlrp3^WT^*, and *hNLRP3^D305N^*mice. *hNLRP3^D305N^* mice demonstrated elevated Nf-L in CSF compared to WT mice (Fig. 1f). Furthermore, *hNLRP3^D305N^* Nf-L increased with age, with levels higher in 13-17-week-old mice compared to 9-10-week-old. This finding suggests that constitutive NLRP3 inflammasome activation promotes progressive neuronal damage.

### *hNLRP3^D305N^* microglia demonstrate distinct reactive transcriptional subpopulations

As resident innate immune cells in the CNS, microglia express the highest levels of *NLRP3* amongst CNS cell types and are critical regulators of neuroinflammation^50,51,42^. Since *hNLRP3^D305N^* mouse brains showed constitutive NLRP3 activity and broad upregulation of proinflammatory signaling proteins, we next sought to elucidate the impact on microglial transcriptional state. We used fluorescence-activated cell sorting (FACS) to capture CD45^+^ cells in the brains of *hNLRP3^WT^*, *mNlrp3^WT^*, and *hNLRP3^D305N^* mice and then conducted scRNA-seq. We used unsupervised clustering to differentiate 14 immune cell types identifiable by their top differentially expressed genes (DEGs) (Supplementary Fig. 3), and isolated microglia *in silico* for further unsupervised clustering analysis to distinguish 7 microglia subclusters (Fig. 2a). *hNLRP3^WT^*and *mNlrp3^WT^* did not show differences in microglia cluster proportions, consistent with most of our results showing no detectable differences between the two control mouse models (Fig. 2b-c). However, the transcriptional signature of the *hNLRP3^D305N^* microglia population was highly differentiated from the WT controls, with changes in abundance observed across subclusters that reflect a general shift from homeostatic microglia towards more reactive states (Fig. 2d). Clusters 0 and 1 had the most significant reductions in *hNLRP3^D305N^* compared to WT and have high expression of canonical homeostatic markers of microglia, such as *Cx3cr1*, *P2ry12*, and *Tmem119*^52^. Cluster 5 exhibited markers of disease-associated microglia (DAM), *Lpl* and *Itgax*^53^, and was also decreased in *hNLRP3^D305N^*, though the overall percentage of this cluster across all genotypes was very low (Fig. 2c). The microglia subclusters enriched in *hNLRP3^D305N^* mice have top DEGs that are associated with microglial reactivity. Cluster 3 constitutes the majority of the *hNLRP3^D305N^* microglial population, and is almost absent in the controls, with top DEGs including *Cd74*, *H2-D1*, *C4b*, *Cxcl13*, and *Apoe* that are associated with various immune responses including antigen presentation, the complement pathway, and lipid homeostasis^54^. Interestingly, these transcripts are broadly expressed across all elevated inflammasome-associated *hNLRP3^D305N^*driven clusters (Fig. 2d) and are most highly expressed in cluster 6, which is also unique to *hNLRP3^D305N^* (Fig. 2c), though far less abundant than cluster 3. Additionally, cluster 6 expresses NLRP3-associated *Il1b*, inflammatory mediator *Spp1*^55^, and *Fn1*, which has been implicated in BBB dysfunction in AD^56^ (Fig. 2d). Finally, the elevation of cluster 4 in *hNLRP3^D305N^* microglia implicates activation of the type I interferon-response, consistent with the pathway’s association with NLRP3^57^. The elevation of these *hNLRP3^D305N^*-associated transcripts, whether broadly expressed or specific to one cluster, can also be detected by microglia pseudobulk analysis (Fig. 2e-f; Supplementary Data 1). Consistent with these findings, the most upregulated gene sets in *hNLRP3^D305N^*microglia compared to *hNLRP3^WT^* are all associated with inflammation (Supplementary Fig. 4; Supplementary Data 2). Taken together, these results demonstrate that constitutive activation of the NLRP3 inflammasome results in broadly reactive and neuroinflammatory microglia, with specific subpopulations (*i.e.,* clusters 3 and 6) that present distinct transcriptional responses compared to pathology-driven microglial subpopulations^53,58,59^.

**Figure 2.**
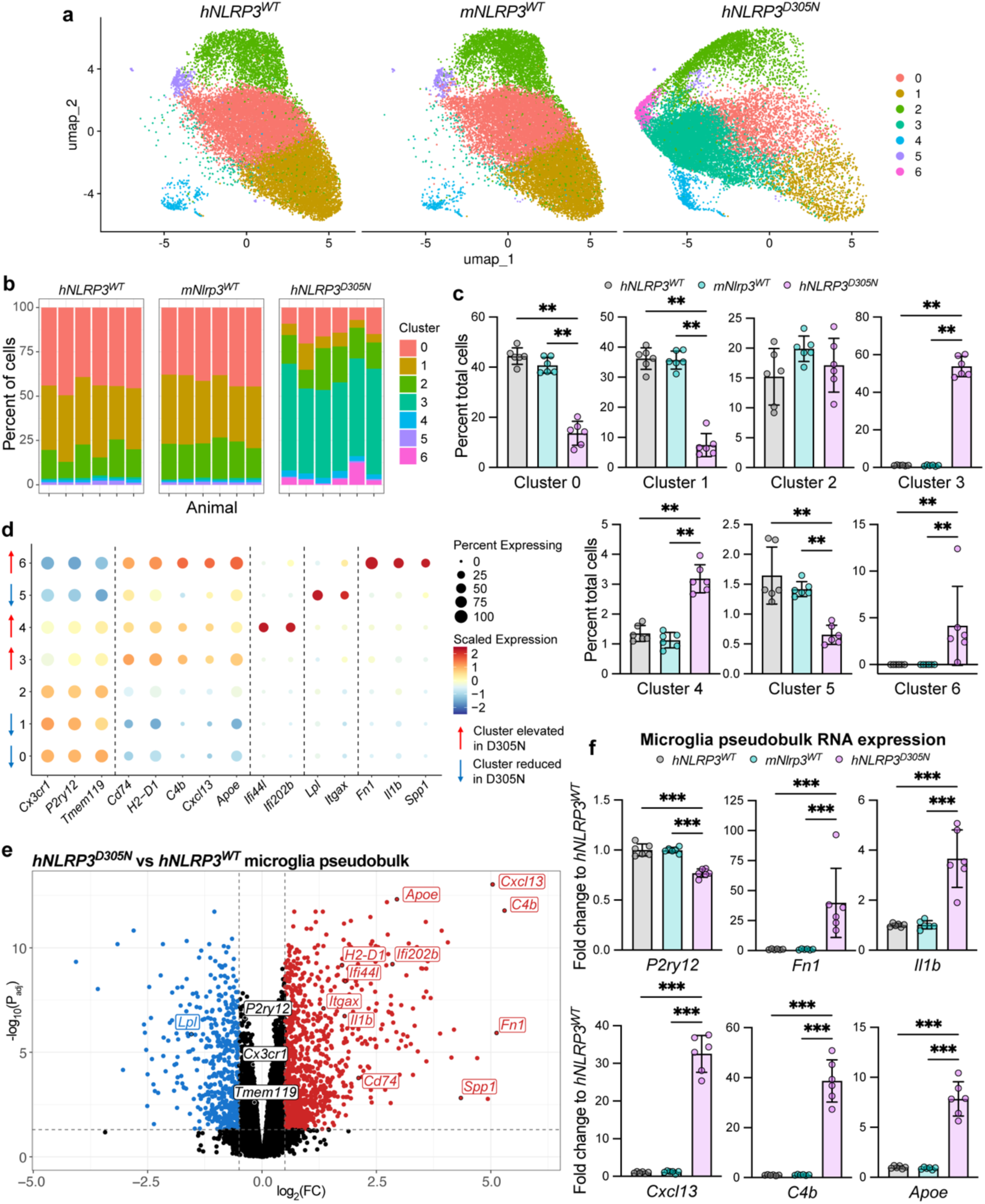
***hNLRP3^D305N^* microglia demonstrate distinct reactive transcriptional subpopulations** Microglia scRNA-seq data from CD45^+^ cells collected via FACS from *hNLRP3^WT^*, *mNlrp3^WT^*, and *hNLRP3^D305N^* mouse brains (Supplementary Fig. 3) (n= 6/group). **(a)** UMAP plots separated by genotype, showing unsupervised clustering into 6 microglia subpopulations. **(b)** Percent of each microglia cluster per animal, separated by genotype. **(c)** Percent of each microglia cluster per genotype. Pairwise Wilcoxon tests across genotype combinations for all clusters with Benjamini-Hochberg multiple testing correction. **(d)** scRNA-seq expression of cluster DEGs and well-described marker genes across microglia clusters. **(e)** Volcano plot of transcripts that are elevated (red) or reduced (blue) in *hNLRP3^D305N^* compared to *hNLRP3^WT^* by microglia pseudobulk RNA-Seq, with transcripts from **(d)** labeled. Only values with p_adj_ < 0.05 and log_2_(FC) > |0.5| are colored. **(f)** Selected DEG top hits from **(d-e)** for *hNLRP3^WT^*, *mNlrp3^WT^*, and *hNLRP3^D305N^*. Lognormal Brown-Forsythe and Welch ANOVA run with Games-Howell’s multiple testing correction for comparisons across genotype. *p_adj_ < 0.05, **p_adj_ < 0.01, ***p_adj_ < 0.001. Data are shown as mean ± SD. Full source data and statistic values in Source Data and Supplementary Data 1.

### Peripheral immune cells traffic to *hNLRP3^D305N^* mouse brains

Previous studies showed peripheral immune cells traffic to the leptomeninges of *hNLRP3^D305N^* mice by qualitative histological visualization of mononuclear and granulocytic cells, and flow cytometry identification of myeloid and lymphoid cells using CD45/CD11b gating^42^. We used our CD45^+^ scRNA-seq dataset to characterize CNS immune cell populations with increased resolution (Supplementary Fig. 3). While brain resident microglia were the most abundant immune cell type, *hNLRP3^D305N^* mice had a significant increase in peripheral immune cell types present compared to the WT control mice (Fig. 3a-b; Supplementary Fig. 5a). The predominant non-microglial immune cell types in *hNLRP3^D305N^* brains were neutrophils (∼9.1%), T cells (∼7.0%), and monocytes (∼6.2%) (Fig. 3c). Regulatory T cells and *Cd4^+^*T cells were the most abundant T cell subsets. This scRNA-seq analysis of CD45^+^ immune cells demonstrates a robust infiltration of various peripheral immune cell populations in the *hNLRP3^D305N^* mouse brain compared to WT, with neutrophils being the most prevalent.

**Figure 3.**
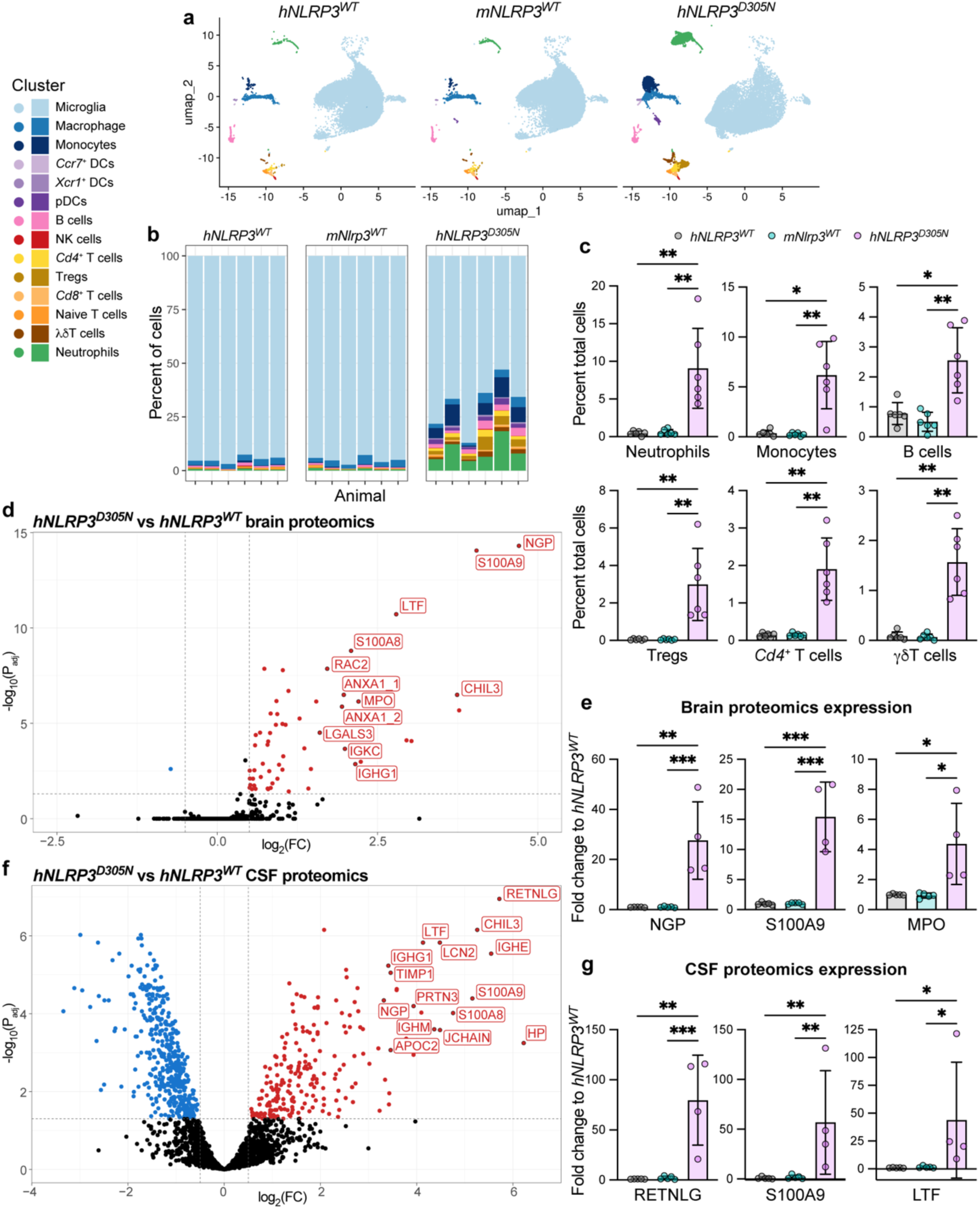
Peripheral immune cells traffic to *hNLRP3^D305N^* mouse brains (a-c) scRNA-seq data collected from CD45^+^ cells collected via FACS from *hNLRP3^WT^*, *mNlrp3^WT^*, and *hNLRP3^D305N^* mouse brains (Supplementary Fig. 3) (n = 6/group). **(a)** UMAP plots colored by cell population (Supplementary Fig. 3) separated by genotype. **(b)** Percent of each cell type per animal, separated by genotype. **(c)** Percent of each cell type for most prevalent cell types. Other cell types shown in Supplementary Fig. 5a. Pairwise Wilcoxon tests across genotype combinations for all cell types with Benjamini-Hochberg multiple testing correction. **(d)** Volcano plot of proteins that are elevated (red) or reduced (blue) in *hNLRP3^D305N^*compared to *hNLRP3^WT^* by untargeted bulk brain proteomics, with top hits labeled (Supplementary Fig. 5c). Only values with p_adj_ < 0.05 and log_2_(FC) > |0.5| are colored (n = 4-5/group). **(e)** Selected bulk brain proteomics top hits from **(d)** for *hNLRP3^WT^*, *mNlrp3^WT^*, and *hNLRP3^D305N^*. Lognormal Brown-Forsythe and Welch ANOVA with Games-Howell’s multiple testing correction for comparisons across genotype (n = 4-5/group). **(f)** Volcano plot of proteins that are elevated (red) or reduced (blue) in *hNLRP3^D305N^* compared to *hNLRP3^WT^* by untargeted CSF proteomics, with top hits labeled (Supplementary Fig. 5d). Only values with p_adj_ < 0.05 and log_2_(FC) > |0.5| are colored (n = 4-5/group). **(g)** Selected CSF proteomics top hits from **(f)** for *hNLRP3^WT^*, *mNlrp3^WT^*, and *hNLRP3^D305N^*. Lognormal Brown-Forsythe and Welch ANOVA with Games-Howell’s multiple testing correction for comparisons across genotype (n = 4-5/group). *p_adj_ < 0.05, **p_adj_ < 0.01, ***p_adj_ < 0.001. Data are shown as mean ± SD. Full source data and statistic values in Source Data and Supplementary Data 3.

Our scRNA-seq analysis served as an opportunity to assess *NLRP3/Nlrp3* expression across these cell types. Previous data indicated *NLRP3/Nlrp3* is predominantly expressed by immune cells in the *hNLRP3^D305N^* and *mNlrp3^WT^* brain compartment, however previous efforts did not differentiate amongst immune cell types^42^.

We detected mouse *Nlrp3* in microglia, macrophages, monocytes, and neutrophils of *mNlrp3^WT^*, consistent with the literature (Supplementary Fig. 5b)^51,60^. Surprisingly, humanized *NLRP3* was not detected in neutrophils, while expression was found in myeloid-lineage cells including microglia, macrophages, and monocytes. There were no apparent differences in *NLRP3* expression in *hNLRP3^WT^* compared to *hNLRP3^D305N^*. Previous characterization of these mouse models showed *NLRP3* was detectable in *hNLRP3^WT^*neutrophils by RT-qPCR (a more sensitive detection method), although expression was lower than *Nlrp3* in *mNlrp3^WT^* neutrophils^41^. Reduced *NLRP3* expression in *hNLRP3^D305N^* neutrophils could attenuate any neutrophil-mediated effects of NLRP3 activation^61^.

We next conducted untargeted proteomics on bulk brain tissue and CSF biofluids from *hNLRP3^WT^*, *mNlrp3^WT^*, and *hNLRP3^D305N^* mice. We observed significantly increased levels of neutrophil-associated proteins, such as neutrophilic granule protein (NGP), S100 calcium-binding protein A9 (S100A9), myeloperoxidase (MPO), resistin-like gamma (RETNLG), and lactotransferrin (LTF), in both the brain and CSF when comparing *hNLRP3^D305N^* mice to the WT mice (Fig. 3d-g; Supplementary Data 3). Other proteins elevated in *hNLRP3^D305N^* mice, selected by highest significant fold change compared to *hNLRP3^WT^*, include proteins that are transcriptionally expressed by monocytes, B cells, and T cells according to our scRNA-seq data (Supplementary Fig. 5c-d). These gene expression patterns suggest a peripheral origin to the presence of these proteins in the bulk brain and/or CSF proteomic data. Altogether these data show that brain infiltration of peripheral immune cells, and particularly neutrophils, is a key pathology in *hNLRP3^D305N^* mice.

Previous studies used scRNA-seq to profile mouse neutrophils from bone marrow, blood, and spleen during homeostasis and after infection^62^. We leveraged this reference atlas to annotate the subpopulation of neutrophils in our dataset (Supplementary Fig. 6a-c). Most of the neutrophils present in *hNLRP3^D305N^*mouse brains belonged to the G5 cluster and specifically the G5c subcluster (Supplementary Fig. 6b), which was identified as a terminally differentiated and apoptotic neutrophil subtype^62^. Among the top upregulated neutrophil-associated proteins in *hNLRP3^D305N^* mice (Supplementary Fig. 5c-d), S100 calcium-binding protein A8 (S100A8)/S100A9^63–65^, lipocalin-2 (LCN2)^66,67^, and LTF^65^ are associated with neutrophil reactivity and neutrophil extracellular trap (NET) release, which are DNA web-like scaffolds associated with proteolytic enzymes that are released by activated neutrophils, often accompanying neutrophil cell death known as lytic NETosis^68,69^. NET-associated proteins^70^ and histones^71^ have been shown to propagate inflammation^72^, induce cytotoxicity^73^, and disrupt BBB integrity^74^. We further confirmed the presence of NET-associated proteins MPO^75^ and S100A9^65^ in *hNLRP3^D305N^* brains by western blot (Supplementary Fig. 6d-e). Altogether, these results suggest that the neutrophils present in *hNLRP3^D305N^*mouse brains are highly mature, reactive, and potentially undergoing increased cell death possibly via NETosis.

### Neutrophils enter the *hNLRP3^D305N^* brain parenchyma

The scRNA-seq and proteomic data demonstrate the presence of neutrophils and other peripheral immune cell types in the brains of *hNLRP3^D305N^*mice, and we have also observed evidence of BBB dysfunction. However, it remains unclear whether these cells are entering into the brain parenchyma or restricted to non-parenchymal brain border regions such as the leptomeninges, a key compartment for CNS immune surveillance^76,77^. Studies in CAPS patients indicate peripheral immune cells accumulate in the leptomeninges^78^, with similar findings in *hNLRP3^D305N^*mice^42^. To examine peripheral immune cell localization, we used IHC in coronal brain sections from *hNLRP3^WT^*and *hNLRP3^D305N^* mice. *mNlrp3^WT^* mice were not used since initial characterizations showed similar phenotypes as *hNLRP3^WT^*(Fig. 1-3). IHC for CD4, a marker well-expressed by helper and regulatory T cells (Supplementary Fig. 3b), revealed that the increase in CNS CD4^+^ T cells was largely restricted to the leptomeninges (Supplementary Fig. 7).

To further evaluate the localization of neutrophils in *hNLRP3^D305N^*mouse brains, we examined the presence of neutrophil-associated proteins lymphocyte antigen 6 complex locus G (Ly6G), S100A9, and MPO by IHC^79^. We detected significantly increased levels of these neutrophil markers in the *hNLRP3^D305N^* mouse brain parenchyma compared to *hNLRP3^WT^* (Fig. 4a-b), matching our proteomic and western blot data (Fig. 3d-g; Supplementary Fig. 6d-e). Notably, both MPO^75^ and S100A9^65^ are associated with inflammatory NET release. Regional quantification confirms correlation among these markers, demonstrating that neutrophil brain parenchymal infiltration is highly region-specific in *hNLRP3^D305N^*, with the highest abundance in the septal nucleus, followed by the corpus callosum, fimbria, hippocampus, and thalamus (Fig. 4c; Supplementary Fig. 8). These brain regions show some proximity to leptomeningeal spaces, which also show high neutrophil abundance as expected (Supplementary Fig. 9). In the case of the hippocampus and thalamus, neutrophil markers were most prevalent near the leptomeningeal borders. These results demonstrate that neutrophils enter the brain parenchyma of *hNLRP3^D305N^* mice in areas near the leptomeninges, an important brain border region for peripheral immune interactions, as well as in more anatomically distinct regions such as the septal nucleus.

**Figure 4.**
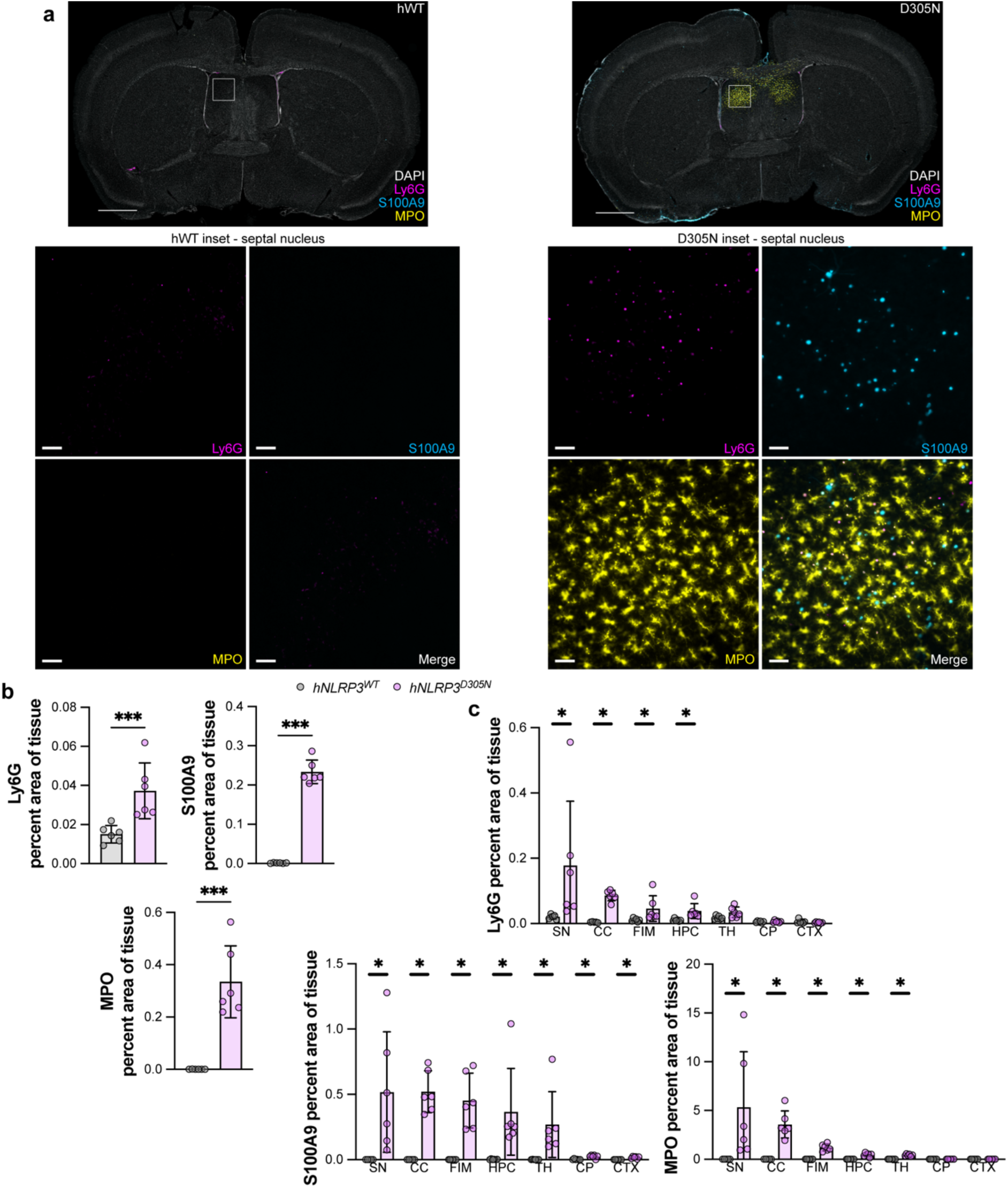
Neutrophils enter the *hNLRP3^D305N^* brain parenchyma. *hNLRP3^WT^* and *hNLRP3^D305N^* whole brain coronal sections were stained for DAPI (white), Ly6G (magenta), S100A9 (cyan), and MPO (yellow) by IHC (n = 6/group). **(a)** Representative widefield images of whole brain coronal sections. Inset is of septal nucleus. Scale bar: whole section, 1000μm; inset, 50μm. **(b)** Quantification of percent of whole tissue stained by Ly6G, S100A9, and MPO. Lognormal Welch two-tailed t-test. **(c)** Quantification of percent of region area stained by Ly6G, S100A9, and MPO. SN, septal nucleus; CC, corpus callosum; FIM, fimbria; HPC, hippocampus; TH, thalamus; CP, caudoputamen; CTX, cortex. Mann-Whitney tests with Holm-Šídák multiple testing correction for genotype comparison in each region. *p_adj_ < 0.05, **p_adj_ < 0.01, ***p_adj_ < 0.001. Data are shown as mean ± SD. Full source data and statistic values in Source Data.

### *hNLRP3^D305N^* microglia demonstrate regional reactivity

Based on the detection of proinflammatory cytokines in the brains of *hNLRP3^D305N^* mice (Fig. 1c), as well as the transcriptional shift in reactive microglial subpopulations (Fig. 2; Supplementary Fig. 4), we sought to visualize canonical microglial markers IBA1 and TMEM119 by IHC in *hNLRP3^D305N^* and *hNLRP3^WT^* coronal whole brain sections (Fig. 5a). We found elevated IBA1 area across the entire brain section in *hNLRP3^D305N^*mice compared to *hNLRP3^WT^* indicative of reactive microglia (Fig. 5b). Although IBA1 staining suggests an increase in the number of microglia, total TMEM119 area was not significantly different between the two genotypes. This may be due to TMEM119 signal being decreased in reactive microglia^80^. Indeed, while IBA1 showed extensive regional increases in *hNLRP3^D305N^*, none of these regions showed a corresponding increase in TMEM119 and the corpus callosum, which had significant neutrophil infiltration in *hNLRP3^D305N^*(Fig. 4c), even showed a decrease in TMEM119 signal compared to WT (Fig. 5c). Notably, all the regions that had significant neutrophil localization in *hNLRP3^D305N^* showed increased IBA1 area compared to *hNLRP3^WT^*, while both regions assessed that did not have signs of neutrophil infiltration in *hNLRP3^D305N^*mice, the caudoputamen and cortex, similarly did not have elevated IBA1 area. Together, these results show that *hNLRP3^D305N^* mouse brains show regional signs of reactive microglia that correlate with neutrophil parenchymal entry.

**Figure 5.**
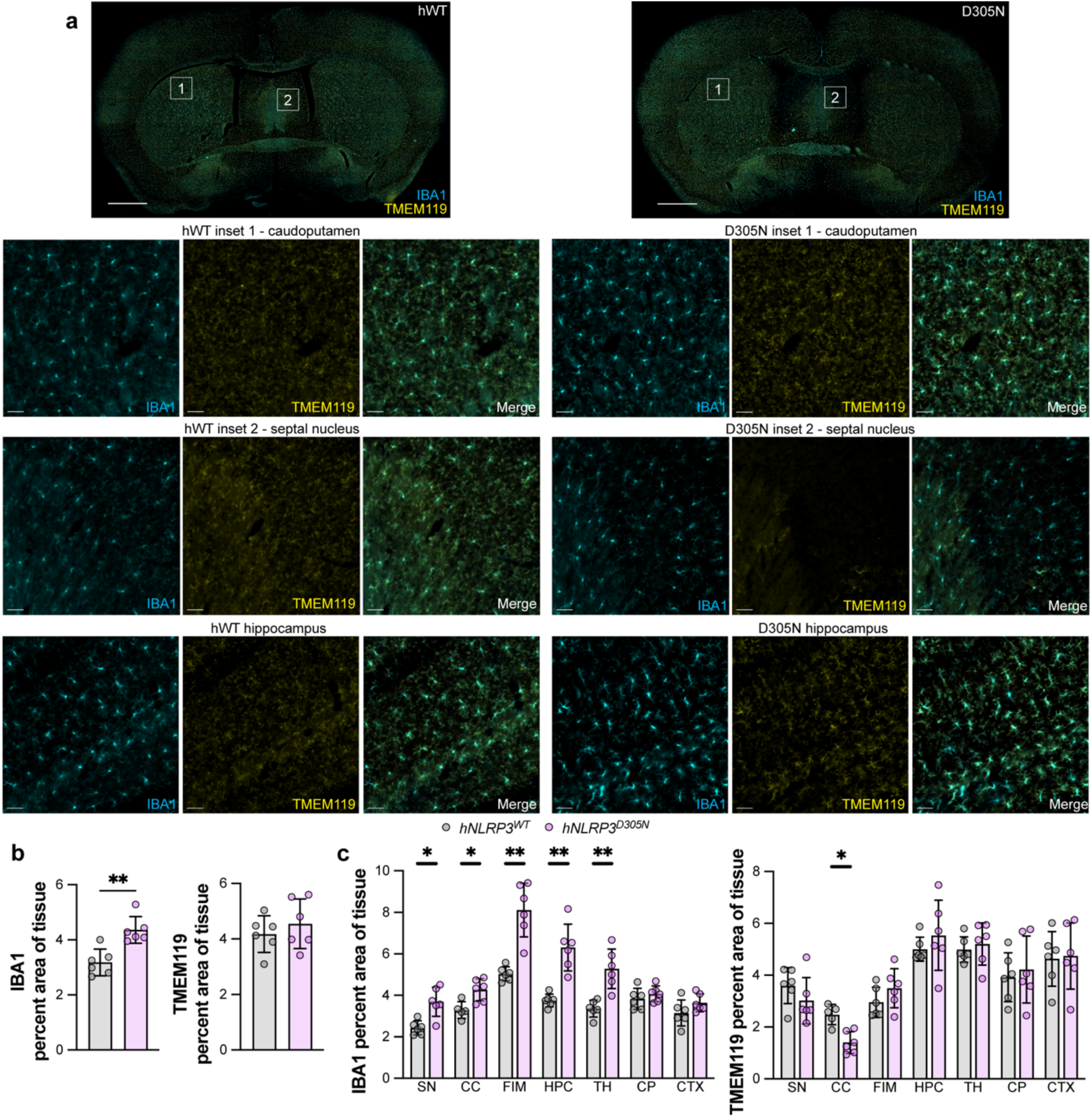
*hNLRP3^D305N^* microglia demonstrate regional reactivity. *hNLRP3^WT^* and *hNLRP3^D305N^* whole brain coronal sections were stained for IBA1 (cyan), and TMEM119 (yellow) by IHC (n = 6/group). **(a)** Representative widefield images of whole brain coronal sections. Inset 1 is of caudoputamen and inset 2 is of septal nucleus. Scale bar: whole section, 1000μm; inset, 50μm. **(b)** Quantification of percent of whole tissue stained by IBA1 and TMEM119. Lognormal Welch two-tailed t-test. **(c)** Quantification of percent of region area stained by IBA1 and TMEM119. SN, septal nucleus; CC, corpus callosum; FIM, fimbria; HPC, hippocampus; TH, thalamus; CP, caudoputamen; CTX, cortex. Lognormal Welch two-tailed t-tests with Holm-Šídák multiple testing correction for genotype comparison in each region. *p_adj_ < 0.05, **p_adj_ < 0.01, ***p_adj_ < 0.001. Data are shown as mean ± SD. Full source data and statistic values in Source Data.

Microglia are highly dynamic and motile cell types that will morphologically adapt depending on functional state^59,81^. Typically, homeostatic microglia present with highly ramified processes to survey the surrounding microenvironment, whereas reactive microglia show relatively retracted processes associated with increased phagocytosis^82,83^. To evaluate microglial morphology in regions of interest and in relation to the presence of invading neutrophils, we used confocal microscopy with LIGHTNING adaptive deconvolution to obtain super-resolution images on the same sections imaged previously^84^ using IBA1^+^ staining to define microglia (Fig. 6a). The increased sensitivity of this imaging approach enabled detection of regional differences in TMEM119 expression, with the average intensity of TMEM119 in microglia being decreased only in the regions of *hNLRP3^D305N^*mouse brains that had clear neutrophil infiltration (Fig. 6b). Furthermore, morphological analysis of microglia similarly demonstrated shifts across multiple metrics, such as decreased mean branch length, increased percent convexity, and decreased mean radius (Fig. 6c), that indicate *hNLRP3^D305N^* microglia were more reactive in mouse brain regions that had neutrophil entry (Fig. 4c). Interestingly, there were modest trends towards reactivity in regions that did not have apparent neutrophil localization, such as a decrease in mean branch length in the caudoputamen of *hNLRP3^D305N^* compared to *hNLRP3^WT^*. Although microglia show morphological heterogeneity across regions, these data indicate that *hNLRP3^D305N^*microglia are more reactive than *hNLRP3^WT^* microglia in areas with neutrophil infiltration, which suggests potential microglia-neutrophil crosstalk.

**Figure 6.**
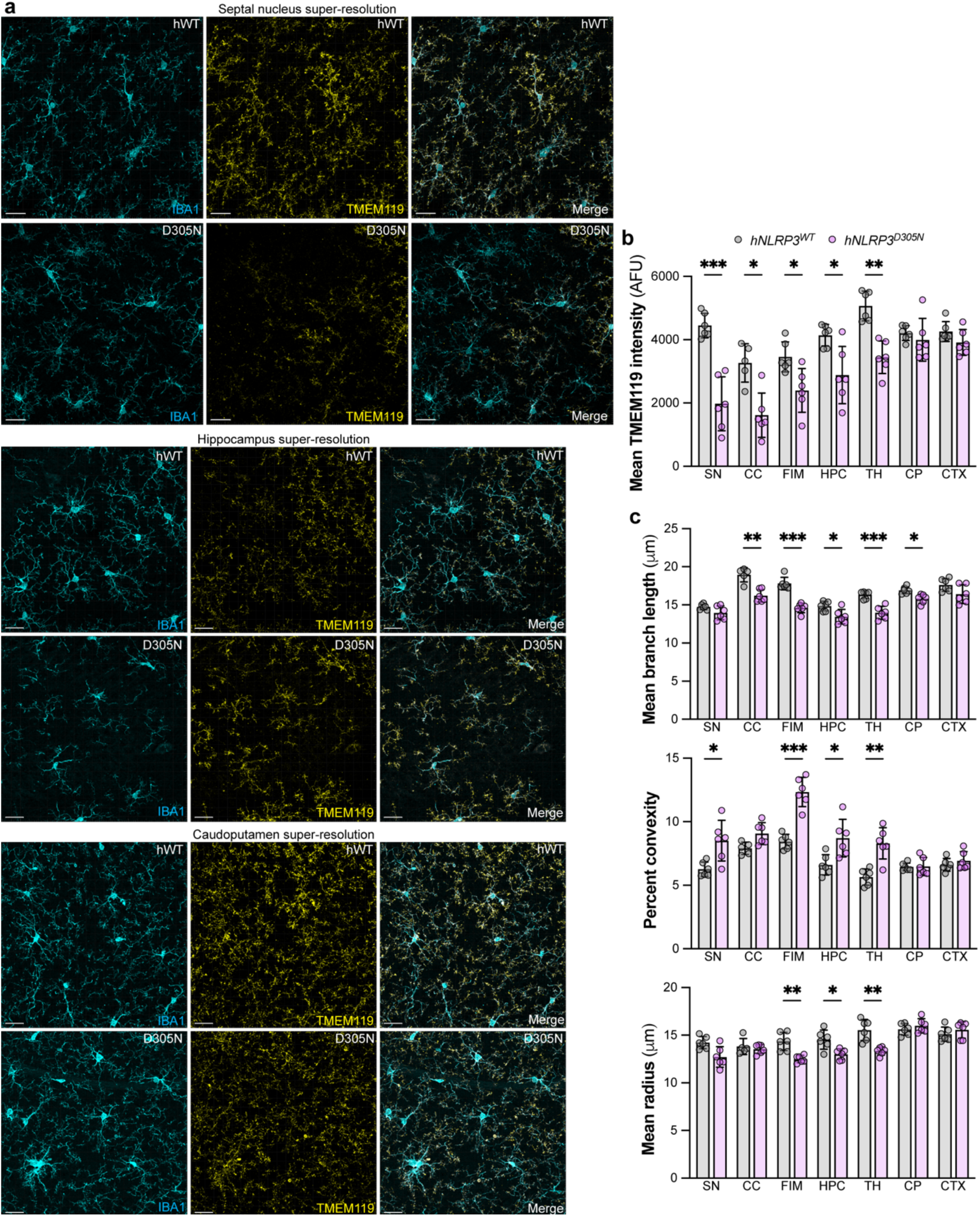
*hNLRP3^D305N^* microglia show morphological features consistent with regional reactivity. *hNLRP3^WT^* and *hNLRP3^D305N^* whole brain coronal sections were stained for IBA1 (cyan), and TMEM119 (yellow) by IHC (n = 6/group). **(a)** Representative confocal super-resolution images of septal nucleus, hippocampus, and caudoputamen. Scale bar: 20μm. **(b)** Quantification of mean TMEM119 intensity per microglia, identified by IBA1 staining in **(a)**, separated by brain region. SN, septal nucleus; CC, corpus callosum; FIM, fimbria; HPC, hippocampus; TH, thalamus; CP, caudoputamen; CTX, cortex. Normal two-tailed t-tests with Holm-Šídák multiple testing correction for genotype comparison in each region. **(c)** Quantification of mean branch length, percent convexity, and mean radius per microglia, identified by IBA1 staining in **(a)**, separated by brain region. SN, septal nucleus; CC, corpus callosum; FIM, fimbria; HPC, hippocampus; TH, thalamus; CP, caudoputamen; CTX, cortex. Normal two-tailed t-tests with Holm-Šídák multiple testing correction for genotype comparison in each region. *p_adj_ < 0.05, **p_adj_ < 0.01, ***p_adj_ < 0.001. Data are shown as mean ± SD. Full source data and statistic values in Source Data.

Astrogliosis is a hallmark feature of neuroinflammation that often occurs alongside activated microglia, where bidirectional signaling between reactive astrocytes and microglia coordinates inflammatory responses in the CNS^85,86^. To investigate potential astrogliosis in *hNLRP3^D305N^* mice, we visualized GFAP, a marker of reactive astrocytes^87^, via IHC (Supplementary Fig. 10a). Consistent with neuroinflammation, GFAP area was significantly elevated in *hNLRP3^D305N^* mouse brains (Supplementary Fig. 10b). Among the regions of interest, quantification of differences in astrogliosis reached significance only in the thalamus and hippocampus, which were both regions that showed robust microglial reactivity (Supplementary Fig. 10c; Fig. 5c; Fig. 6b-c). These results demonstrate that *hNLRP3^D305N^* mouse brains have astrogliosis that shows partial regional correlation with reactive microglia and neutrophil infiltration.

### Microglia and neutrophils show direct interactions in *hNLRP3^D305N^*mouse brains

Since *hNLRP3^D305N^* mouse brains show regional neutrophil infiltration and those regions are also associated with changes in microglial reactivity, we next investigated whether neutrophils and microglia physically interact in these regions. NET release by neutrophils has been shown to activate inflammatory pathways in microglia^88^ and microglia have even been described as engulfing neutrophils in models of cerebral ischemia^89,90^. To investigate potential cell-cell interactions in *hNLRP3^D305N^* mouse brains, we stained coronal whole brain sections with antibodies against IBA1, Ly6G, and MPO (Fig. 7a) and found ∼35% of Ly6G^+^ neutrophil staining across all brain regions assessed colocalized with IBA1^+^ microglia, suggesting that these cell types are directly interacting (Fig. 7b). Notably, most of the Ly6G-IBA1 colocalization was also MPO^+^, further suggesting that the neutrophils are activated and potentially undergoing NETosis. Surprisingly, although MPO is associated with neutrophils^75^, the total percentage of MPO signal that colocalized with Ly6G was very low. Most of the MPO signal detected either colocalized with IBA1 or did not colocalize with IBA1 or Ly6G. This result may suggest that MPO is being secreted or released by neutrophils in the brain parenchyma, indicative of NET release and/or NETosis^91^, which has also been associated with neutrophil transmigration across the BBB^68^. To better visualize the potential interactions between neutrophils and microglia, we used confocal super-resolution microscopy on the same sections (Fig. 7c), which showed instances of Ly6G^+^ spherical neutrophils being engulfed by IBA1^+^ microglia processes (Fig. 7d). These results suggest that Ly6G^+^ neutrophils enter the *hNLRP3^D305N^* brain parenchyma, where they likely degranulate, release MPO, and undergo some degree of phagocytosis by activated microglia, potentially inducing further microglial reactivity. IBA1, Ly6G, and MPO show partial colocalization suggestive of microglia-neutrophil interactions.

**Figure 7.**
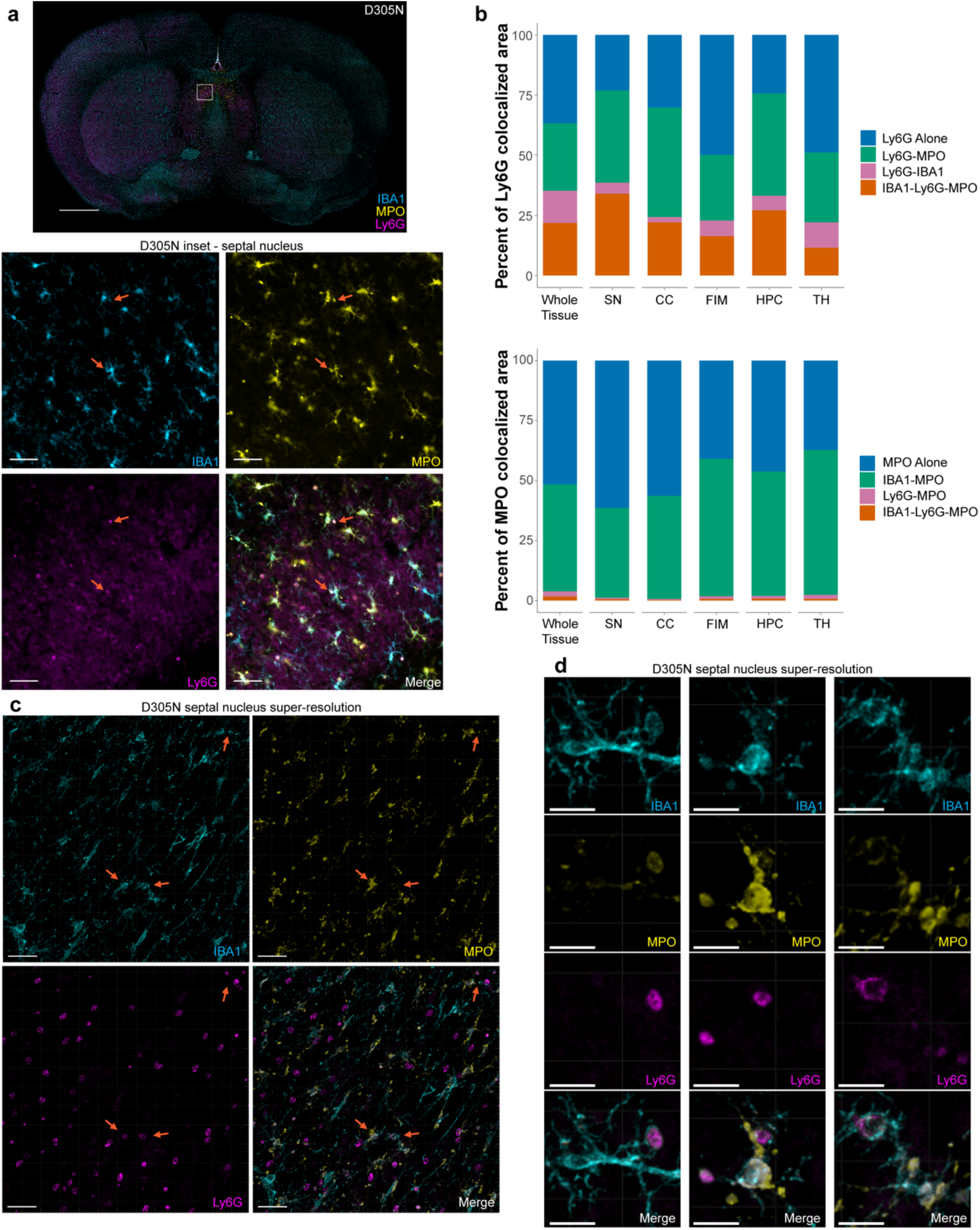
Microglia and neutrophils show direct interactions in *hNLRP3^D305N^* mouse brains. *hNLRP3^D305N^* whole brain coronal sections were stained for IBA1 (cyan), MPO (yellow), and Ly6G (magenta) by IHC (n = 6). **(a)** Representative widefield image of whole brain coronal section. Inset is of septal nucleus. Arrows point to areas with colocalization between all three markers. Scale bar: whole section, 1000μm; inset, 50μm. **(b)** Percent of Ly6G or MPO signal that colocalizes with other IHC markers, separated by brain region. SN, septal nucleus; CC, corpus callosum; FIM, fimbria; HPC, hippocampus; TH, thalamus. **(c)** Representative confocal super-resolution images of septal nucleus. Arrows point to areas with colocalization between all three markers. Scale bar: 30μm. **(d)** Representative confocal super-resolution images of septal nucleus. Scale bar: 10μm. Full source data and statistic values in Source Data.

## Discussion

The NLRP3 inflammasome is emerging as a druggable pathway in neurodegenerative diseases, where evidence points to its involvement both upstream and downstream of various neuropathologies^1–3^. In this study, we demonstrate in the *hNLRP3^D305N^* GoF mouse model that chronic activation of the NLRP3 inflammasome is sufficient to drive sterile neuroinflammation, BBB permeability, peripheral immune cell recruitment, and progressive neuronal damage in the absence of neuropathological aggregates or any other external stimuli. Specifically, we show that constitutive NLRP3 activation in GoF human mutation carrying mice leads to extensive proinflammatory cytokine and chemokine production in the brain, broad transcriptional shifts towards distinct reactive microglial subpopulations, diverse peripheral immune cell CNS infiltration identified by scRNA-seq and bulk proteomics, robust region-specific neutrophil entry into the brain parenchyma, regional morphological changes in microglia consistent with microglial hyperreactivity, astrogliosis, and increased Nf-L in the CSF suggestive of neuronal damage. Our findings provide evidence that NLRP3 activation is sufficient to initiate a feed-forward inflammatory cascade capable of inducing BBB permeability and neuronal damage, while revealing neutrophil-microglia immune cell interactions.

Microglia show the highest *NLRP3* expression of all CNS cell types^50,51,42^ and transcriptomic analysis in *hNLRP3^D305N^* mice revealed the emergence of a unique, highly transcriptionally reactive subpopulation (cluster 6). Notably, cluster 6 had the highest expression of many inflammatory and disease-associated transcripts that were broadly upregulated among *hNLRP3^D305N^*microglia, such as *Apoe*, *C4b*, and *Cxcl13*, while also uniquely expressing NLRP3-associated *Il1b*. Interestingly, other top DEGs in cluster 6 include *Spp1* and *Fn1*, which both encode extracellular matrix glycoproteins that are often upregulated under inflammatory conditions and implicated in immune cell trafficking^92,93^. An *Spp1^+^* and *Fn1^+^* profibrotic macrophage subpopulation was recently identified in a murine model of myocardial infarction^94^, suggesting that microglia-mediated fibrosis may play a role in the CNS phenotypes present in *hNLRP3^D305N^*mice. The degree to which these transcriptional changes in microglia may be influencing neutrophil recruitment remains to be determined. Concurrently, the microglia in regions of neutrophil parenchymal infiltration may be undergoing further transcriptional remodeling through their interactions with neutrophils, leading to functional changes that could have a major impact on the neuroinflammatory state of the brain, BBB homeostasis, and ultimately neurodegeneration.

A CNS-penetrant small molecule NLRP3 inhibitor was recently shown to reduce peripheral immune cell infiltration into the brain and leptomeninges of *hNLRP3^D305N^* mice by flow cytometry and qualitative histology, while a peripherally restricted NLRP3 inhibitor did not^43^. Given the central role of microglia in regulating neuroinflammation^95^ and the marked increase in a broad panel of proinflammatory cytokines and chemokines in *hNLRP3^D305N^* brains, we speculate that reactive *NLRP3* GoF microglia may act as key effectors driving the production of factors that recruit neutrophils into the brain parenchyma. Candidate signaling proteins elevated in *hNLRP3^D305N^* brains that are associated with neutrophil activation and recruitment include IL-1β^96–99^, G-CSF^100–102^, IL-6^103–105^, KC^40,46,106,107^, IL-17^108–110^, and LIX^106,111,112^, although we cannot exclude other candidates from the literature^113^ not detected in this study. Indeed, because the humanized model used has low *NLRP3* expression in neutrophils, this suggests that the *hNLRP3^D305N^* mutation may be driving neutrophil phenotypes, including brain infiltration, primarily via non-cell-autonomous mechanisms. Interestingly, our findings diverge from previous research that showed a neutrophil-specific *Nlrp3* GoF mutation was sufficient to drive neutrophil migration to the brain and BBB permeability^40^. The discrepancy here may be due to the humanization of *NLRP3*, as functional differences have been reported between mouse and human *NLRP3* promoters and GoF mutations^42,114^. It is also possible that neutrophil NLRP3 activation can cause their migration to the brain, but that this is not the primary mechanism present in *hNLRP3^D305N^* mice, where region-specific neutrophil recruitment into the CNS may be primarily driven by local, CNS-intrinsic chemotactic signals derived from microglia or perhaps border-associated macrophages with NLRP3 pathway activation.

One of the most striking results of our study was the profound recruitment of multiple peripheral immune cell types into the brains of *hNLRP3^D305N^*mice. Although an influx of peripheral immune cells has been reported previously in the leptomeninges of *hNLRP3^D305N^* mice under baseline conditions^42^, we examined immune cell populations with single cell resolution and demonstrated significant entry of neutrophils into the brain parenchyma by IHC. The leptomeninges have emerged as an important immunological surveillance niche for the brain in health and disease, including neurodegenerative diseases^76,115,77,116^. The regional patterns of parenchymal neutrophil infiltration and broad peripheral immune cell trafficking into the leptomeninges in *hNLRP3^D305N^* mice suggest that meningeal brain entry routes and signaling pathways may be relevant in this model. Notably, a subset of neutrophil proteins detected in the CNS (*e.g.,* S100A9 and MPO) by untargeted proteomics, western blot, and IHC are associated with reactive neutrophils and NETs^65,75^. Furthermore, comparing the transcriptional profile of the neutrophils present in *hNLRP3^D305N^* mouse brains to previously reported subpopulations from the literature^62^ linked the most prevalent neutrophil subpopulations with reactive profiles, including apoptotic pathways. These findings support previous studies that have described a synergistic feed-forward relationship between inflammasome activation and NET release in varied disease contexts^117^, and the reduced levels of *NLRP3* in *hNLRP3^D305N^* neutrophils suggest a non-cell-autonomous association. Since neutrophils and NETs can propagate inflammation^72^, induce cytotoxicity^73^, and disrupt BBB integrity^74^ in other settings, neutrophil transmigration and activation, or even lytic NETosis, may contribute to BBB dysfunction and neurodegeneration-associated phenotypes in *hNLRP3^D305N^* mice.

Using confocal super-resolution microscopy, we observed direct physical neutrophil-microglia interactions consistent with microglial engulfment of neutrophils. Microglial phagocytosis of neutrophils has been reported previously in rodent models of cerebral ischemia^89,90^. Moreover, neutrophils have been shown to activate microglial inflammatory pathways in models of stroke and traumatic brain injury^88,118,119^. Notably, our data showed that MPO colocalizes with IBA1^+^ microglia, indicating microglial uptake of this protein either via neutrophil secretion, possibly in the form of NETs, or during microglial phagocytosis of neutrophils. While microglia appeared generally more reactive in the *hNLRP3^D305N^* mouse brains by IHC, consistent with NLRP3 activity, microglia in regions of neutrophil entry showed changes in morphology and protein expression indicative of greater reactivity. This finding shows consistency with the microglia transcriptomic data, which also demonstrates broad inflammatory shifts (cluster 3) in conjunction with the emergence of a specific hyperreactive subpopulation (cluster 6). These results suggest that microglia-neutrophil bidirectional crosstalk may be critical to the neuropathologies present in *hNLRP3^D305N^* mice driven by constitutive NLRP3 activation.

Importantly, this *hNLRP3^D305N^* mouse model demonstrated *NLRP3* expression and pathway activation in both CNS and peripheral myeloid cells, along with inflammation in both the CNS and periphery^41^. Thus, any CNS phenotypes observed, such as neuroinflammation, changes in BBB permeability, and neuronal damage, could potentially be driven in part by systemic inflammation and peripheral NLRP3 activity. Indeed, systemic LPS administration has been shown to induce similar phenotypes in previous studies^46,120^. While the literature suggests that myeloid cells, and microglia in the CNS, are likely the primary initiators of the NLRP3 inflammasome activity in the *hNLRP3^D305N^*mouse model^42^, we did not investigate *NLRP3* expression in other potential CNS-relevant cell types such as endothelial cells^121^, astrocytes^122^, or neurons^30^. Establishing cell type-specific *NLRP3* expression in this GoF model could help distinguish the contributions of microglial vs peripheral immune cell NLRP3 pathway activation on the CNS phenotypes we describe herein.

Although NLRP3 has been implicated in neurodegenerative diseases, this study demonstrates that genetic NLRP3 activation appears sufficient to drive neuronal damage. Nf-L is emerging as one of the key biomarkers for neurodegeneration in both pre-clinical^48,123^ and clinical settings^49,124,125^, however, it is generally unclear whether its detection in biofluids reflects frank neuronal loss, axonal degeneration, or a combination of both occurring in CNS tissue, among other possible influences^126^. Our study does not provide any direct evidence for frank neuronal loss or axonal degeneration at a gross level, but we cannot exclude selective neuronal vulnerability escaping detection using pan-neuronal or axonal markers. Our results also demonstrate that *hNLRP3^D305N^*mice have broad neuroinflammation, increased BBB permeability, and robust peripheral immune cell recruitment to the brain. Neutrophils are shown to be a key cell type in this model that appear to directly interact with microglia in the brain parenchyma, suggesting that this interaction may be crucial towards the CNS phenotypes described. Overall, our findings establish NLRP3 inflammasome activation as a central and sufficient driver of neuroinflammatory and neurodegeneration-associated processes, highlighting its potential as a therapeutic target while underscoring the need to further dissect the cellular and mechanistic interplay between peripheral immune infiltration and CNS-resident responses that may be relevant in neurodegenerative diseases and CAPS. Finally, our study suggests CNS or CSF neutrophil-associated biomarkers potentially define a signature of chronic NLRP3 activation that may inform selection of patient subpopulations and pathway endpoints in clinical settings, given the emergence of investigational CNS-penetrant NLRP3 inhibitors.

## Methods

### Antibodies, reagents, and supplies

All catalog numbers for reagents used in methods can be found in Supplementary Table.

### Mouse handling

Generation of *hNLRP3^WT^* and *hNLRP3^D305N^*129S6 mice was described previously^41^. *mNlrp3^WT^* mice were littermates of *hNLRP3^D305N^* mice, bred using *hNLRP3^D305N^*/*mNlrp3^WT^*heterozygotes. All mice characterized in this study were homozygous for *mNlrp3^WT^*, *hNLRP3^WT^*, or *hNLRP3^D305N^*. Mice were bred and maintained by The Jackson Laboratory and shipped to Denali Therapeutics no less than one week before initiating any experimental procedures. All mouse procedures adhered to regulations and protocols approved by Denali Therapeutics Institutional Animal Care and Use Committee (IACUC). Mice were housed under a 12-hour light/dark cycle and group housed when possible. Sex was as evenly split as possible between male and female for all groups in all studies. Age and sex were as evenly distributed among groups within a study as possible.

Mouse study cohort overview:

| Figure | Groups (n) | Age range (wk) |
| --- | --- | --- |
| 1a-c, 1f, S1d | *hNLRP3<sup>WT</sup>* (6), *mNlrp3<sup>WT</sup>* (9), *hNLRP3<sup>D305N</sup>* (6) | 9 - 10 |
| 1a-c, 1f, S1d | *hNLRP3<sup>WT</sup>* (10), *mNlrp3<sup>WT</sup>* (13), *hNLRP3<sup>D305N</sup>* (13) | 13 - 17 |
| 1d-e | *hNLRP3<sup>WT</sup>* (7), *mNlrp3<sup>WT</sup>* (6), *hNLRP3<sup>D305N</sup>* (7) | 9 - 13 |
| S1a-b, S2a-b, S2d-e, S6d-e | *hNLRP3<sup>WT</sup>* (8), *hNLRP3<sup>D305N</sup>* (8) | 9 - 10 |
| S1c | *hNLRP3<sup>WT</sup>* (8), *mNlrp3<sup>WT</sup>* (12), *hNLRP3<sup>D305N</sup>* (15) | 18 - 23 |
| S2c | *hNLRP3<sup>WT</sup>* (8), *hNLRP3<sup>D305N</sup>* (6) | 21 - 24 |
| 2, S3, S4, 3, S5,<br>S6a-c | *hNLRP3<sup>WT</sup>* (6), *mNlrp3<sup>WT</sup>* (6), *hNLRP3<sup>D305N</sup>* (6) | 12 - 15 |
| S7, 4, S9, 5, 6, S10,<br>7 | *hNLRP3<sup>WT</sup>* (6), *hNLRP3<sup>D305N</sup>* (6) | 19 - 20 |

### Mouse tissue collection

For all tissue collection, animals are anesthetized via intraperitoneal injection of 2.5% avertin. For plasma, blood is collected via cardiac puncture into EDTA-coated tubes, which were then centrifuged at 12,700 rpm for 7 min at 4°C before the top layer was collected. Cerebrospinal fluid (CSF) was extracted vs cisterna magna puncture and spun down at 12,700 rpm for 7 min at 4°C before the supernatant was collected. Fluids were frozen on dry ice and stored at-80°C. Mice were perfused with ice-cold PBS at a rate of 5 mL/min for 3-5 min. For biochemical analysis, tissues were collected and weighed before freezing on dry ice and storing at-80°C. For immunohistochemistry (IHC), brains were drop-fixed in 4% paraformaldehyde (PFA) in PBS for 24 hrs at 4°C with gentle agitation before transferring to PBS with 0.01% sodium azide at 4°C. If animals were only used for IHC (*i.e.,* in cohort with coronal whole-brain images), animals were perfused with 4% PFA in PBS at room temperature at a rate of 2.5 mL/min for 15 min after PBS perfusion and before drop-fixing.

### Tissue homogenization for protein assays

Weighed tissue samples were submerged in RIPA buffer with cOmplete protease inhibitor and PhosStop phosphatase inhibitor (100 μL of buffer per 10 mg of tissue). Samples were then homogenized using a 3mm tungsten carbide bead, shaken using the QIAGEN TissueLyzer II (2×3 min at 27 Hz). Samples were centrifuged at 18,000 g for 30 min at 4°C, then supernatant was aliquoted for analysis. BCA was used to quantify protein levels. Samples were stored at-80°C.

### IL-1β MSD

MSD V-PLEX Mouse IL-1β Kit was used to assess IL-1β levels. Plasma and tissue homogenate were diluted 2x in provided sample diluent and manufacturer instructions were followed using provided supplies and MSD MESO SECTOR S 600 instrument. Tissue IL-1β concentrations were normalized by total protein assessed by BCA.

### IL-1RA and IL-18 ELISA

R&D Systems Mouse IL-1ra/IL-1F3 Quantikine ELISA Kit was used to assess IL-1RA levels and MBL Mouse IL-18 ELISA Kit was used to assess IL-18 levels. For IL-1RA analysis, plasma and tissue homogenate were processed neat and manufacturer instructions were followed using provided supplies. For IL-18 analysis, plasma and tissue homogenate were diluted 5x in provided assay diluent and manufacturer instructions were followed using provided supplies. Plates were read using Agilent Technologies BioTek Synergy Neo2, running Gen5 software. Tissue concentrations were normalized by total protein assessed by BCA.

### Western blot

Tissue homogenate was normalized after BCA assessment then denatured in NuPAGE Sample Reducing Agent and NuPAGE LDS Sample Buffer at 70°C for 10 min. 20 μg per sample was loaded into NuPAGE 4 to 12%, Bis-Tris, 1.0 mm Protein Gels in NuPAGE MES SDS Running Buffer. Proteins were subsequently transferred to Bio-Rad Trans-Blot Turbo Midi 0.2 μM Nitrocellulose Transfer Packs with a Bio-Rad Trans-Blot Turbo Transfer System. Membranes were washed three times with TBST, blocked in Rockland Blocking Buffer for Fluorescent Western Blotting for 1 hr at room temperature on a rocking platform, and incubated with primary antibodies diluted in Rockland Blocking Buffer for 16-20 hrs at 4°C on a rocking platform. The following day, membranes were washed three times with TBST, incubated in secondary antibodies diluted in Rockland Blocking Buffer at room temperature on a rocking platform protected from light, washed three times with TBST, and visualized using an LI-COR Biosciences Odyssey CLx Imager. Immunoblots were analyzed using Image Studio 5.2. Quantification was reported as band signal after median background subtraction. If the value was negative due to extremely low true signal, this value was replaced with 0. Supplementary Fig. 1a-b, Supplementary Fig. 2a-b, and Supplementary Fig. 6d-e use the same western blot.

### IgG MSD

A generic MSD-based platform sandwich electrochemiluminescence immunoassay was used to quantify total concentration of mouse IgG antibodies in mouse plasma and brain lysate. Thermo Scientific Blocker Casein was used as assay diluent and added to an MSD GOLD 96-well Small Spot Streptavidin SECTOR Plate at 150 μL/well for approximately 1 hr. Following plate blocking and a plate wash step, a biotinylated goat anti-mouse IgG polyclonal antibody at a working concentration of 1 μg/mL in assay diluent was added to coat the assay plate at 30 μL/well, and was incubated for 1-2 hrs. Following the plate coating and a plate wash step, test samples were added to the assay plate at 25 μL/well, and incubated for 1-2 hrs. Following the sample incubation and a plate wash step, a secondary ruthenylated (SULFO-TAG) goat anti-mouse IgG polyclonal antibody at a working solution of 1 μg/mL in assay diluent was added to the assay plate at 25 μL/well, and was incubated for approximately 1 hr. 1x MSD Read Buffer T was then added to generate the electrochemiluminescence (ECL) assay signal, which was read using a MSD MESO SECTOR S 600 instrument.

All assay reaction steps were performed at ambient temperature and with shaking on a plate shaker at a speed of ∼650 rpm (where appropriate). Any unknown plasma samples with projected concentrations outside the assay standard curve range would be subject to additional sample pre-dilutions with assay diluent for quantifiable measurements. Sample ECLU signals generated in the assay were subsequently processed into sample concentrations by back-calculating off the assay calibration standard (CS) curve. This assay had a dynamic CS range of 0.0305 – 200 ng/mL in Assay Diluent with a total of 9 standard points (serially-diluted at 1:3) plus a blank matrix sample. The assay CS curve was fitted with a weighted four-parameter non-linear logistic regression using MSD Discovery Workbench 4.0 for use in calculating concentrations for unknown/test samples.

### Cytokine analysis

Brain homogenate was normalized to 5 mg/mL total protein after BCA assessment then frozen on dry ice. Samples were shipped and processed by Eve Technologies (Canada) for the Mouse Cytokine/Chemokine 44-Plex Discovery Assay (MDF44).

### CSF neurofilament light chain Simoa

Quanterix Simoa NF-light Advantage kit was used to assess neurofilament light chain levels as described previously^123,48^. CSF was diluted 100x in provided sample diluent and manufacturer instructions were followed using provided supplies and Quanterix HD-X instrument.

### Brain immunohistochemistry

Fixed mouse brains (see Mouse tissue collection) were coronally sectioned at NeuroScience Associates using their MultiBrain sectioning service. Briefly, brains were trimmed and mounted in a single gelatin block, then coronally sectioned at a thickness of 40μm. Gelatin sheets with embedded brain sections were then stored in antigen preservation solution (50% ethylene glycol + 1% PVP in PBS) until staining. Brain sections were placed under broad-spectrum LED lights for 48-72 hrs in PBS at 4°C to reduce non-specific fluorescence. Sections were equilibrized to room temperature, then incubated in Antigen Unmasking Solution, Tris-Based at 80°C for 30 min. Sections were washed 3 times with PBST (0.05% Tween in PBS) before incubating in blocking buffer (1% BSA + 0.1% fish gelatin + 0.5% Triton X-100 + 0.1% sodium azide in PBS) for 1 hr at room temperature with gentle agitation. Sections were then transferred to antibody dilution solution (1% BSA + 0.3% Triton X-100 + 0.1% sodium azide in PBS) with primary antibodies and gently agitated at 4°C for 16-20 hrs. Sections were then washed 3 times with PBST before transferring to antibody dilution solution with secondary antibodies for 1-2 hrs at room temperature with gentle agitation protected from light. Samples were then washed for 8 min with PBS with 0.5 μg/mL DAPI in PBST before washing 2 more times with PBST. Finally, sections were mounted onto 2’’ x 3’’ slides with ProLong Glass Antifade Mountant and allowed to dry overnight at room temperature.

### Immunohistochemistry widefield image collection and analysis

Widefield images were acquired using a Zeiss Axioscan.Z1 slide scanner using a 20x/0.8 NA air objective and images were processed using Zeiss ZEN 3.12 software. Automated shading correction and stitching in Zen was then used to produce single complete images for each section. Identical imaging settings and post-processing methods were used to collect images from all tissue sections from the experiment in a single imaging run. For IgG and albumin, quantification was reported as total staining intensity normalized by tissue area. For Ly6G, MPO, S100A9, IBA1, and TMEM119, positively stained objects were identified using local intensity threshold or Intellesis trainable segmentation, then filtered with size selection to remove tiny imaging artifacts. Quantification was reported as total positive area normalized by tissue area. For colocalization quantification, object masks for each channel were overlaid over the microscopy image to identify colocalization area. The Allen Mouse Brain Common Coordinate Framework v3 (CCFv3)^127,128^ was used as a reference to manually annotate brain regions of interest on slide scanner images of whole coronal mouse brain sections. Sections were manually inspected and found to match the atlas at roughly three depths. Seven total brain regions were annotated: the septal nucleus, caudoputamen, corpus callosum (medial corpus callosum between apex of cingulum bundles), the cortex (layers 5/6 of the primary somatosensory area), fimbria, thalamus, and hippocampus. See Supplementary Fig. 8 for examples of regions outlined.

### Immunohistochemistry confocal super-resolution image collection and analysis

Confocal images were acquired using a Leica Microsystems SP8 scanning confocal microscope using a 25x/0.95 NA water objective and a z-step of 0.5 μm, collecting multiple fields of view per-animal, per-brain region. Images were post-processed with Leica Lightning deconvolution in adaptive mode using Leica LAS X (v3.5.7) and then each file was imported into Oxford Instruments Imaris (v10.0) for visualization. To analyze the microglia morphology, the IBA1 channel was used to segment microglia and assess morphology following a modification of a previously described protocol^129^. To speed processing, images were first downsampled 2x in x and y using the transform.downscale_local_mean function in scikit-image (v0.22)^130^. Non-specific background was removed by smoothing the image with a gaussian filter with an ∼15 μm isotropic sigma then subtracting the filtered image from the original image and setting all negative values to 0 using the ndimage.gaussian_filter function in scipy (v1.13)^131^. Images were then thresholded with a manually determined threshold followed by removal of connected components smaller than ∼8.4 μm^3^, then filling of mask holes smaller than ∼8.4 μm^3^, then a binary closing (voxel-wise dilation followed by erosion) using the functions in the morphology module of scikit-image. The filtered mask was first split into connected components using the ndimage.label function in scipy, then masks larger than ∼1500 μm^3^ were further split into sub-1500 μm^3^ components using k-means clustering in scikit-learn (v1.4)^132^ with k set to the smallest value that would result in clusters smaller than the volume threshold and each voxel in the mask assigned to the nearest cluster center. Each segment was skeletonized using the morphology.skeletonize function in scikit-image. Finally, per-segment statistics were calculated for mean TMEM119 intensity within the mask, mean length of uninterrupted degree 1 and 2 voxel segments in the skeleton (mean branch length), mean distance from the center of mass to each voxel in the segment (mean radius), and the ratio of the volume of each segment to the volume of the convex hull of the segment (percent convexity) as calculated using the ConvexHull class in scipy. These metrics were then averaged across all replicate images per-animal, per-brain region.

### Single cell RNA-sequencing sample and library preparation

18 mice (6 per genotype: *mNlrp3^WT^, hNLRP3^WT^, hNLRP3^D305N^*) were processed for single-cell RNA-sequencing (scRNA-seq) in two batches of nine animals each. Whole brains (excluding olfactory bulb and cerebellum) were dissociated using the Miltenyi Adult Mouse Brain Dissociation Kit with modifications. Enzyme Mix 1 was supplemented with anisomycin (2 μM), actinomycin D (5 μM), and hyaluronidase Type I-S (5,600 U/mL) to enhance extracellular matrix digestion and inhibit transcription and translation.

Single-cell suspensions were co-labeled with SyTox and CD45 for 30 min on ice. After washing, 0.5–1 × 10⁶ cells per sample were labeled with unique 10x Genomics Cell Multiplexing Oligos (CMO) per the manufacturer’s protocol, substituting PBS with NbActiv-1 + 1% BSA. Each batch of nine CMO-labeled samples was counted by hemocytometer, pooled at equal cell numbers, FACS-sorted for CD45⁺ immune cells, and loaded onto a Chromium Chip G following the Chromium NextGEMS Single Cell 3′ v3.1 Cell Multiplexing Rev B protocol (CG000388), targeting ∼3,000 immune cells per animal. Following reverse transcription, GEM cleanup, and cDNA amplification, products were separated into 3′ gene expression (GEX) and CMO fractions via dual-sided size selection. Five microliters of CMO cDNA were PCR-amplified to add Illumina indices, and one-quarter of the GEX cDNA was used for library preparation. GEX libraries were pooled equimolarly and quality-checked on an Illumina MiSeq to estimate cell recovery. Based on these results, GEX and CMO libraries were re-pooled and sequenced on an Illumina NovaSeq S2 (28 × 10 × 10 × 90 cycles) at SeqMatic (Fremont, CA), targeting 50,000 reads per cell for mRNA and 5,000 reads per cell for CMO demultiplexing.

### Single cell RNA-sequencing analysis of all immune cells

Raw sequencing reads were aligned to the mouse reference genome (mm10) using Cell Ranger multi (10x Genomics, v7.0.1). Downstream analyses were performed in R^133^ (v4.4.1) using Seurat (v5.3.0)^134,135^.

Cells with fewer than 500 detected features, more than 4,500 features, or greater than 5% mitochondrial transcripts were excluded. Gene expression counts were log10-transformed and normalized using the “*NormalizeData”* function. Highly variable genes (n = 2,000) were identified using “*FindVariableFeatures”* with the “vst” method. Data were scaled using default parameters in “*ScaleData”*, and principal component analysis (PCA) was performed with RunPCA. Data from multiplexed sequencing pools were integrated in principal component (PC) space using canonical correlation analysis (CCA), treating each pool as a batch. Uniform Manifold Approximation and Projection (UMAP) embeddings were generated from the first 30 principal components using “*RunUMAP”*. The same PCs were used for graph-based clustering with “*FindNeighbors”* and “*FindClusters”* (resolution = 0.5) using the Louvain algorithm. Doublets were identified and removed using scDblFinder (v1.20.2)^136^. Broad cell type annotations were assigned based on marker genes identified with “*FindAllMarkers*” and established canonical markers. Contaminating non-immune brain cell types (*e.g.,* neurons and oligodendrocytes) were excluded from further analysis. To obtain higher-resolution annotations, NK/T cells and monocyte/macrophage/dendritic cell populations were subset, reclustered independently, and annotated using FindAllMarkers in conjunction with known marker genes. These refined annotations were subsequently transferred back to the full dataset. The percentage of each cell type expressing *NLRP3* (human) or *Nlrp3* (mouse) was calculated from log-normalized RNA expression values. Cells were classified as “expressing” if the log-normalized expression value was greater than 0. To estimate the percentage of expressing cells included in visualizations, the proportion of cells with normalized expression > 0 of the gene of interest was computed within each cell type/cluster, pooling all cells regardless of sample. For all scRNA-seq analyses described (global immune cell analysis, microglia, and neutrophils), cell type or cell state abundances were calculated per sample as the percentage of all cells passing quality control within the full dataset or within the relevant subset. Statistical significance was assessed using Wilcoxon rank-sum tests for all pairwise comparisons. P values were adjusted for multiple testing using the Benjamini–Hochberg method, and results were considered statistically significant at an adjusted *P* value < 0.05. All visualizations were generated using ggplot2 (v4.0.0)^137^. The top 15 proteins with significant fold changes from the brain and CSF proteomics analyses were mapped to their mouse orthologs and evaluated for cell type–specific expression using scRNA-seq data. Duplicate entries, proteins lacking a mouse ortholog, and genes without appreciable expression in the scRNA-seq dataset were excluded. The remaining genes (10 from brain dataset, 12 from CSF) were ordered by hierarchical clustering using “*hclust”* from the stats package (v4.4.1), and dendrogram data were extracted using “*dendro_data”* from ggdendro (v0.2.0)^138^. The dendrogram and dot plot were combined using “*plot_grid”* from cowplot (v1.1.3)^139^.

### Single cell RNA-sequencing analysis of microglia

For microglia-specific analyses, the dataset was subset to microglial cells. Clustering was performed across multiple resolutions (0–1.5), and cluster stability was evaluated using clustree (v0.5.1)^140^. A resolution of 0.2 was selected for downstream analyses. Differentially expressed genes (DEGs) between clusters were identified using “*FindAllMarkers*”. For pseudobulk analysis, microglia were aggregated by sample ID and coarse cell type using “*aggregateAcrossCells*” from scuttle (v1.16.0)^141^. Differential gene expression analyses comparing *hNLRP3^D305N^* to *hNLRP3^WT^* genotypes were performed using limma/voom (v3.62.2)^142^. Pseudobulked expression data were normalized using the trimmed mean of M values (TMM) method implemented in “*calcNormFactors”* using edgeR (v4.4.2)^143^. Log-transformed counts per million (logCPM) were computed using “*cpm”* with a prior count of 2. For genes of interest, fold change for each sample was calculated relative to the mean logCPM expression of *hNLRP3^WT^* mice. Gene set enrichment analysis (GSEA) was conducted using fgsea (v1.32.4)^144,145^ on the *hNLRP3^D305N^* versus *hNLRP3^WT^* comparison. Hallmark gene sets were obtained from the Molecular Signatures Database (MSigDB) using msigdbr (v25.1.1)^146,147^.

### Single cell RNA-sequencing analysis of neutrophils

For neutrophil-specific analyses, cells annotated as neutrophils were subset for downstream state characterization and reference mapping. A previously published neutrophil reference atlas^62^, encompassing differentiation states from early bone marrow precursors to mature circulating neutrophils, was used for cell state annotation. Author-provided cluster annotations were retained. Low-quality and contaminating cells were removed according to the methodology described in the original publication. Batch correction was performed in principal component space using harmony (v1.2.4)^148,149^, treating each sample as a separate batch. Neutrophil subpopulation annotations were transferred from the reference atlas using Seurat’s label transfer workflow^150^. Transfer anchors were identified using “*FindTransferAnchors*”, and predicted labels were assigned with “*TransferData*”.

### Proteomics data generation and statistical analysis

Quantitative proteomic analysis of mouse brain tissue lysates was performed at BGI Americas (San Jose, CA) using tandem mass tag (TMT)–based multiplexed labeling. The overall workflow followed established Denali proteomics methods previously described for TMT-based tissue proteomics^151^, with adaptations for batch structure and fractionation depth. Tissue samples were lysed in SDS-containing buffer, and protein concentrations were determined prior to digestion. Proteins were reduced with dithiothreitol (DTT), alkylated with iodoacetamide (IAM), and digested with Trypsin/Lys-C. Following digestion, peptides were desalted using C18 solid-phase extraction and dried under vacuum. Peptides were reconstituted in triethylammonium bicarbonate (TEAB) buffer and labeled with isobaric TMT reagents according to the manufacturer’s instructions. Sample channel assignments followed the accompanying annotation file. Labeling efficiency was verified by a preliminary LC–MS/MS label-check run prior to pooling. TMT-labeled peptides were combined in equal amounts and fractionated by high-pH reversed-phase chromatography into 12 fractions per batch. Each fraction was analyzed by nano-LC coupled to a high-resolution Orbitrap mass spectrometer (*e.g.,* Q Exactive HF-X or Orbitrap Eclipse) using data-dependent acquisition with MS3-based reporter-ion quantification to reduce ratio compression. Peptide identification, protein inference, and reporter-ion quantification were performed by BGI Americas using their optimized in-house analysis pipeline. Protein-level quantitative data were provided to Denali Therapeutics for downstream normalization and statistical analysis.

Label-free proteomics analysis of mouse CSF was conducted at Denali Therapeutics using a data-independent acquisition (DIA) workflow on a Bruker timsTOF Pro 2 mass spectrometer operated in dia-PASEF mode. CSF samples were processed by in-solution digestion, including protein denaturation, reduction with dithiothreitol (DTT), alkylation with iodoacetamide (IAM), and enzymatic digestion with trypsin or Trypsin/Lys-C, followed by C18 desalting. These preparation steps are consistent with core DIA proteomics workflows previously reported by Denali Therapeutics using the same platform^152^, with modifications appropriate for low-protein CSF samples. Peptides were separated by reversed-phase chromatography using an Agilent 1290 system configured for split-flow nano-liquid chromatography and coupled to the timsTOF Pro 2. DIA acquisition employed dia-PASEF with a precursor mass range of approximately 300–1400 m/z, an ion-mobility range of ∼0.7–1.4 1/K0, and a TIMS ramp time of ∼100 ms. Collision energy was applied using an ion-mobility–dependent scheme. DIA isolation windows were optimized using py_diAID. Raw data were processed in Spectronaut v19 using a comprehensive spectral library generated from project-matched samples and reference proteomes. Peptide and protein identifications were controlled at a 1% false discovery rate. Protein-level quantitative matrices were generated following normalization and quality-control filtering to remove low-confidence features and proteins with excessive missing values. For both brain and CSF proteomics data, the *hNLRP3^D305N^*versus *hNLRP3^WT^* comparison was evaluated. Differential abundance analysis was performed using limma with array weights estimated by arrayWeights. Proteins were tested against a minimum log2 fold change threshold of log2(1.1) using the treat function, and p-values were adjusted for multiple comparisons using the Benjamini-Hochberg method^142^.

## Statistical analyses

All individual data points shown represent distinct animals. As much as possible, samples for an experiment in a study were run together to minimize experimental variability. For biological data that make scientific sense to calculate a ratio (*e.g., NLRP3* expression is 0.5x compared to control) and values ≤ 0 do not exist, we assumed a lognormal distribution since biological data is often affected by multiplicative processes that result in a lognormal distribution (or at least more often lognormal than normal)^153^. Thus, for most comparisons we use a lognormal t-test or ANOVA. For ease of visual interpretation, we graph the arithmetic mean ± SD on a linear scale. When we have values of 0, we either use a normal ANOVA (*e.g.,* Fig. 1e) or a non-parametric Mann-Whitney test (*e.g.,* Fig. 4c). We made no assumptions about equal SDs between groups, hence the Brown-Forsythe and Welch tests. Unless otherwise noted, we corrected for multiple comparisons using the recommended test by GraphPad Prism 10 (version 10.6.0), which we used often for visualization and statistical analysis. Sex was considered as a biological variable and assessed during initial characterization experiments. Since there was no apparent sex effect on the phenotypes identified for genotype, we did not characterize the impact of sex for subsequent studies. Full source data and statistic values are available in Source Data.

## Data Availability

Full source data and statistic values are available in the Source Data provided with this paper. scRNA-seq data is available via GEO (GSE328214). Proteomics data will be made available via the MassIVE repository upon publication.

## Code Availability

Analysis code will be made available in Zenodo upon publication.

## Acknowledgements

We thank Chandrani Chakraborty for assistance in initial exploratory experiments, Gabrielly Lunkes de Melo for assistance with scRNA-seq preparation, Audrey G. Reeves for assistance with the IgG MSD, and Hoda Safari Yazd and Sonnet S. Davis for assistance with LC-MS. We thank Annie Arguello, Melodi Bowman, and Amy Wing-Sze Leung for support and training for mouse surgical procedures.

## Author contributions

LLS and GDP conceptualized and designed this study. LLS, RC, IB, CC, and HNN performed mouse surgical procedures. LLS, RC, and DH performed biochemical characterization experiments. LLS performed IHC staining and imaging. LLS, DJ, and JCD performed image analysis. LLS, CH, LS, and EWS performed mouse brain scRNA-seq. SMG and DT performed scRNA-seq analyses. NSG and JHS performed proteomic data generation and analysis. EWS, TS, RGT, and PJL provided feedback and suggestions throughout. LLS, SMG, SVA, KMM, and GDP wrote the manuscript. All authors edited the manuscript and provided comments.

---

**Supplementary Figure 1.**
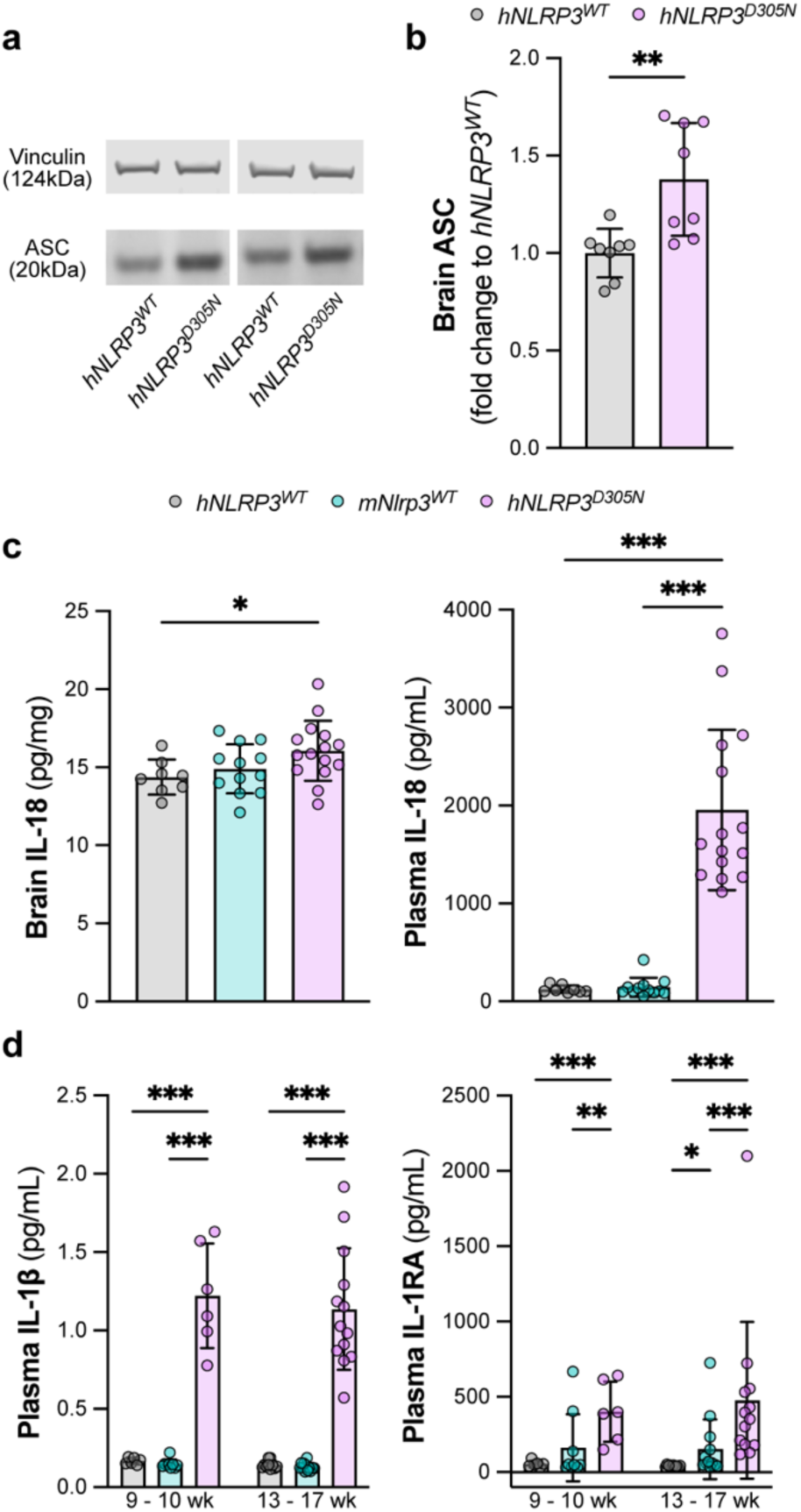
The NLRP3 inflammasome pathway is constitutively active in the brain and peripherally in *hNLRP3^D305N^* mice (a-b) Representative images and quantification of western blot for ASC in bulk mouse brains for *hNLRP3^WT^*and *hNLRP3^D305N^*. Vinculin used as loading control. Lognormal Welch two-tailed t-test (n = 8/group). **(c)** IL-18 ELISA quantification of bulk mouse brains and plasma for *hNLRP3^WT^*, *mNlrp3^WT^*, and *hNLRP3^D305N^*. Lognormal Brown-Forsythe and Welch ANOVA with Tukey multiple testing correction for comparisons across genotype (n = 8-15/group). **(d)** IL-1β MSD and IL-1RA ELISA quantification of plasma samples for *hNLRP3^WT^*, *mNlrp3^WT^*, and *hNLRP3^D305N^*. Lognormal two-way ANOVA with Tukey multiple testing correction for comparisons across genotype and age group (n = 6-13/group). *p_adj_ < 0.05, **p_adj_ < 0.01, ***p_adj_ < 0.001. Data are shown as mean ± SD. Full source data and statistic values in Source Data.

**Supplementary Figure 2.**
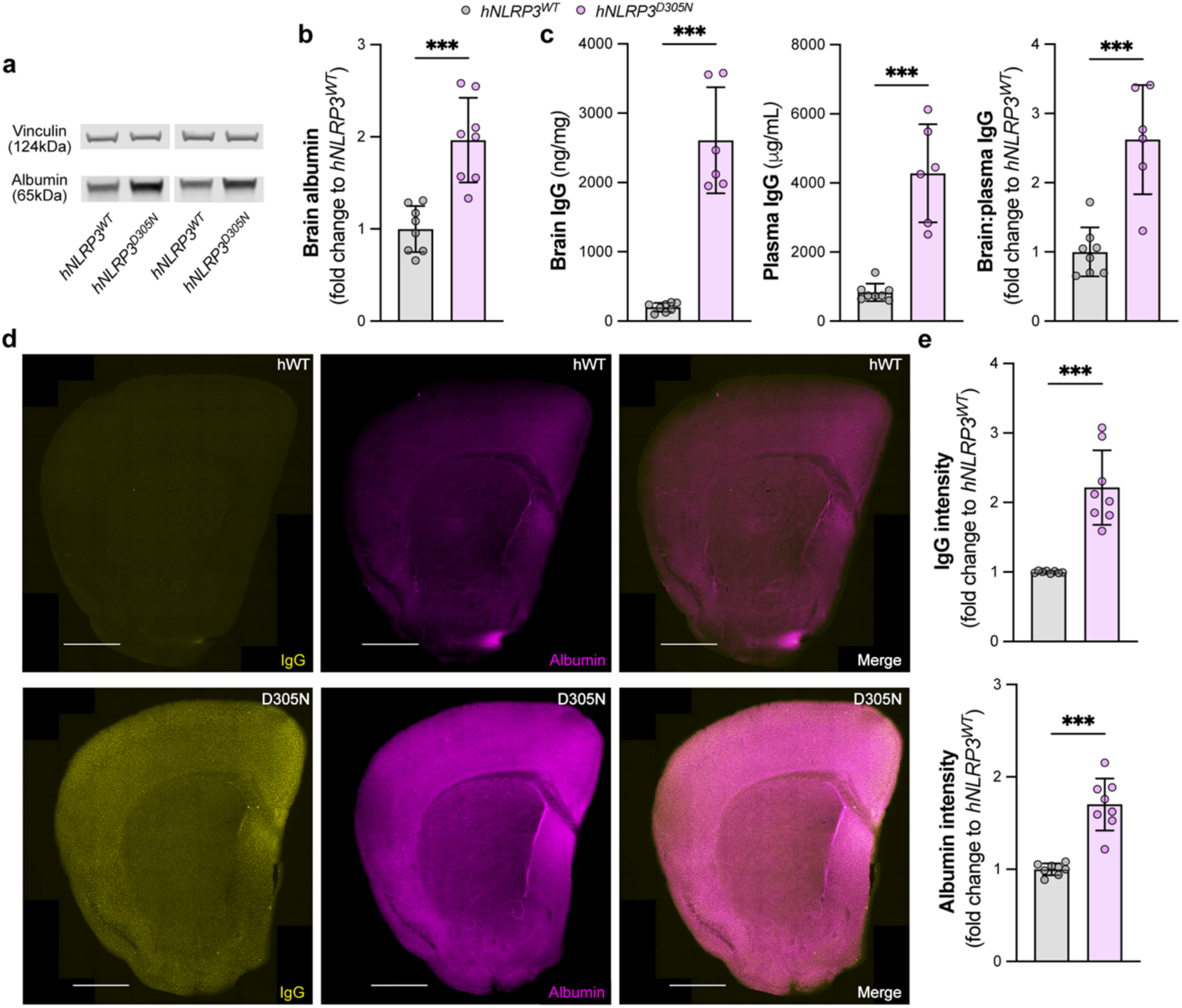
Blood-associated proteins present in *hNLRP3^D305N^* mouse brains are indicative of blood-brain barrier permeability (a-b) Representative images and quantification of western blot for albumin in bulk mouse brains for *hNLRP3^WT^* and *hNLRP3^D305N^*. Vinculin used as loading control. Lognormal Welch two-tailed t-test (n = 8/group). **(c)** IgG MSD quantification of bulk brain, plasma, and brain:plasma ratio for *hNLRP3^WT^* and *hNLRP3^D305N^*. Lognormal Welch two-tailed t-test (n = 6-8/group). **(d-e)** Representative widefield images and quantification of IgG (yellow) and albumin (purple) IHC in mouse coronal brain hemisphere sections for *hNLRP3^WT^* and *hNLRP3^D305N^*. Lognormal Welch two-tailed t-test. Scale bar: 1000μm (n = 8/group). *p_adj_ < 0.05, **p_adj_ < 0.01, ***p_adj_ < 0.001. Data are shown as mean ± SD. Full source data and statistic values in Source Data.

**Supplementary Figure 3.**
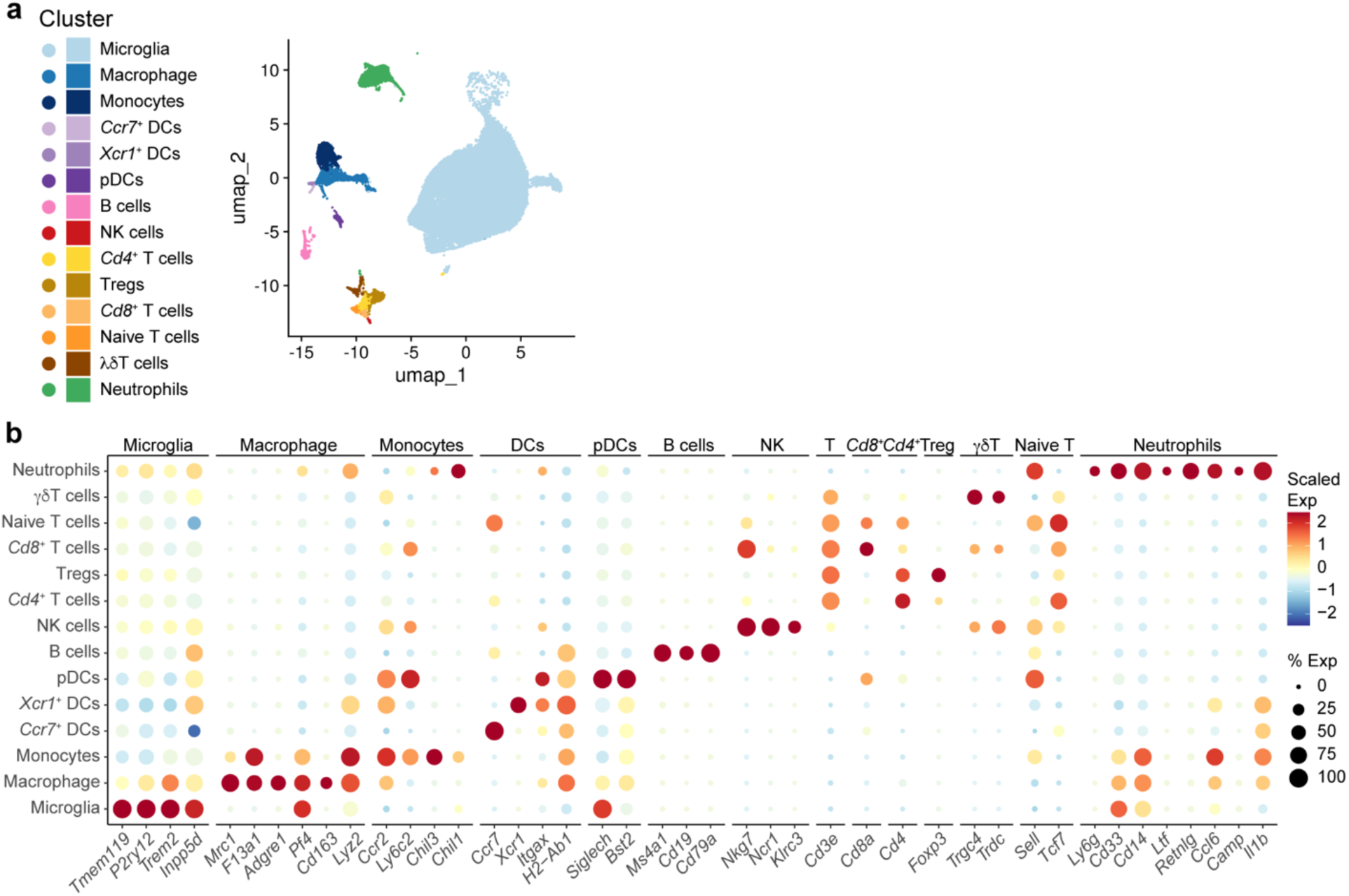
scRNA-seq cell type clustering of CNS CD45^+^ immune cells. scRNA-seq data collected from CD45^+^ cells collected via FACS from *hNLRP3^WT^*, *mNlrp3^WT^*, and *hNLRP3^D305N^* mouse brains (n = 6/group). **(a)** UMAP plot showing unsupervised clustering into 14 immune cell populations that were identified by top DEGs. **(b)** scRNA-seq expression of cluster DEGs and well-described marker genes across clusters, showing cell type identification. Full source data and statistic values in Source Data.

**Supplementary Figure 4.**
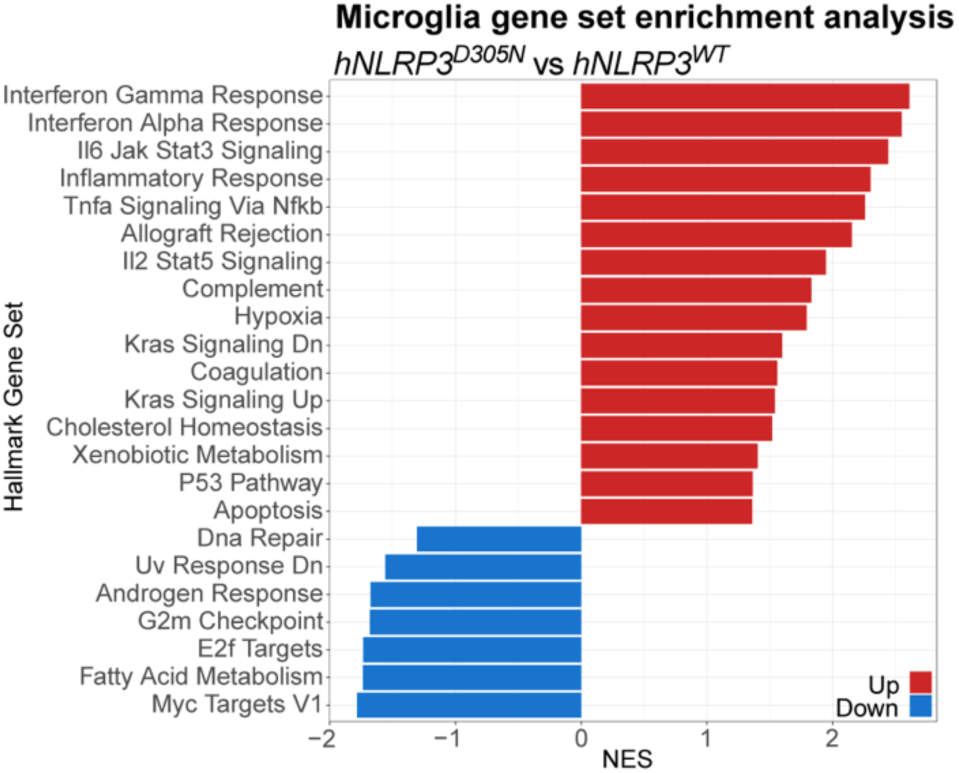
*hNLRP3^D305N^*microglia show upregulated reactive transcriptional pathways. Microglia gene set enrichment analysis of microglia pseudobulk RNA-Seq data (Fig. 2e), showing Hallmark Gene Set pathways that are elevated (red) or reduced (blue) in *hNLRP3^D305N^* compared to *hNLRP3^WT^*as measured by normalized enrichment score (NES) (n = 6/group). Pathways reaching p_adj_ < 0.1 are shown. Full source data and statistic values in Source Data and Supplementary Data 2.

**Supplementary Figure 5.**
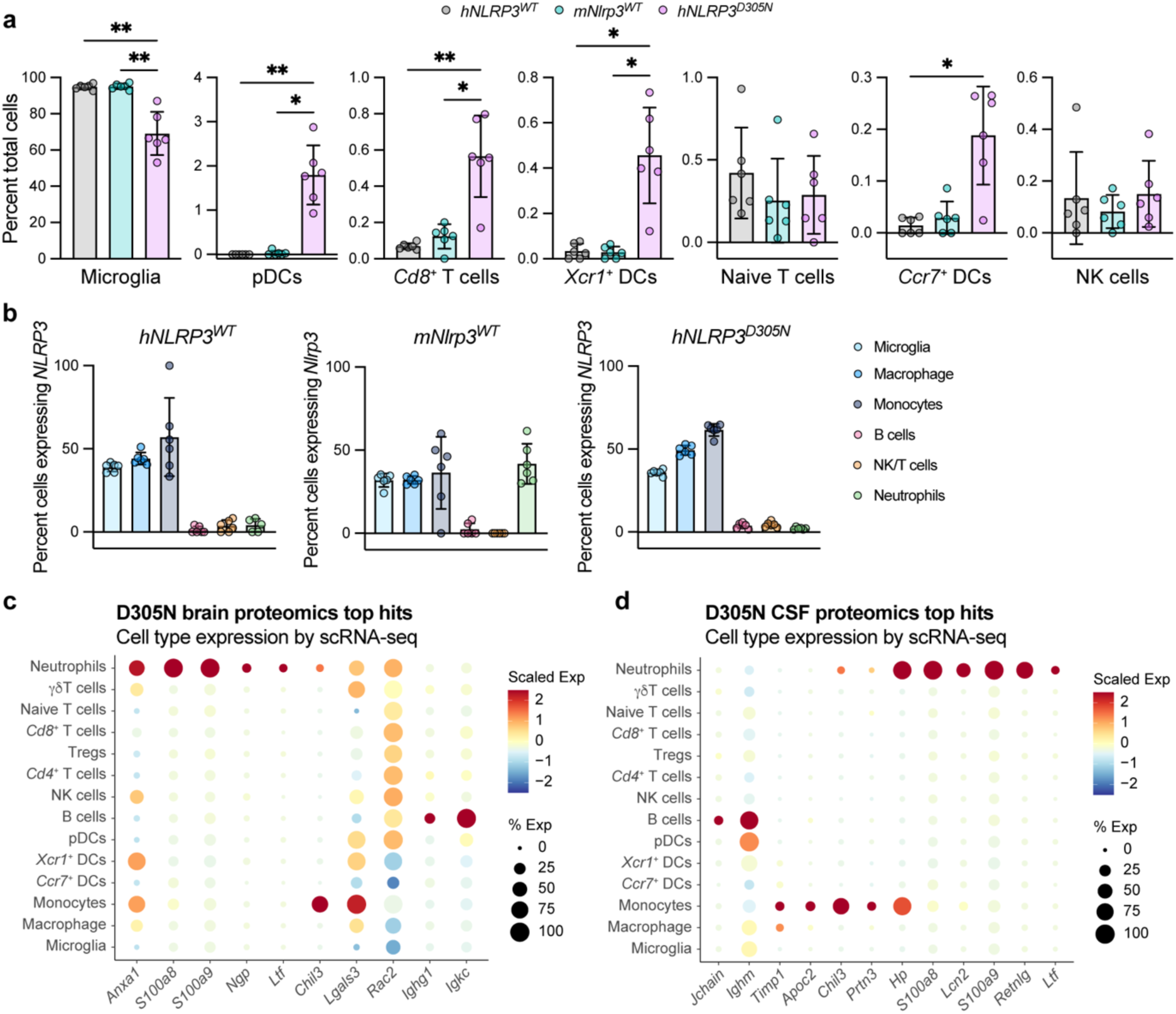
Peripheral immune cells traffic to *hNLRP3^D305N^* mouse brains. scRNA-seq data collected from CD45^+^ cells collected via FACS from *hNLRP3^WT^*, *mNlrp3^WT^*, and *hNLRP3^D305N^* mouse brains (Supplementary Fig. 3) (n = 6/group). **(a)** Percent of each cell type for profiled cell types not shown in Fig. 3c, from scRNA-seq dataset. Pairwise Wilcoxon tests across genotype combinations for all cell types with Benjamini-Hochberg multiple testing correction. **(b)** Percent of cells in a cell type that express *NLRP3/Nlrp3,* shown for each genotype, from scRNA-seq dataset. **(c-d)** scRNA-seq expression of top proteomics hits from Fig. 3d bulk brain and Fig. 3f CSF across cell types identified by scRNA-seq. For each sample type, the 20 most elevated proteins in *hNLRP3^D305N^* compared to *hNLRP3^WT^*that had p_adj_ < 0.05 were probed against the scRNA-seq dataset for matching transcripts that were detected in > 5% of any cell type. *p_adj_ < 0.05, **p_adj_ < 0.01, ***p_adj_ < 0.001. Data are shown as mean ± SD. Full source data and statistic values in Source Data.

**Supplementary Figure 6.**
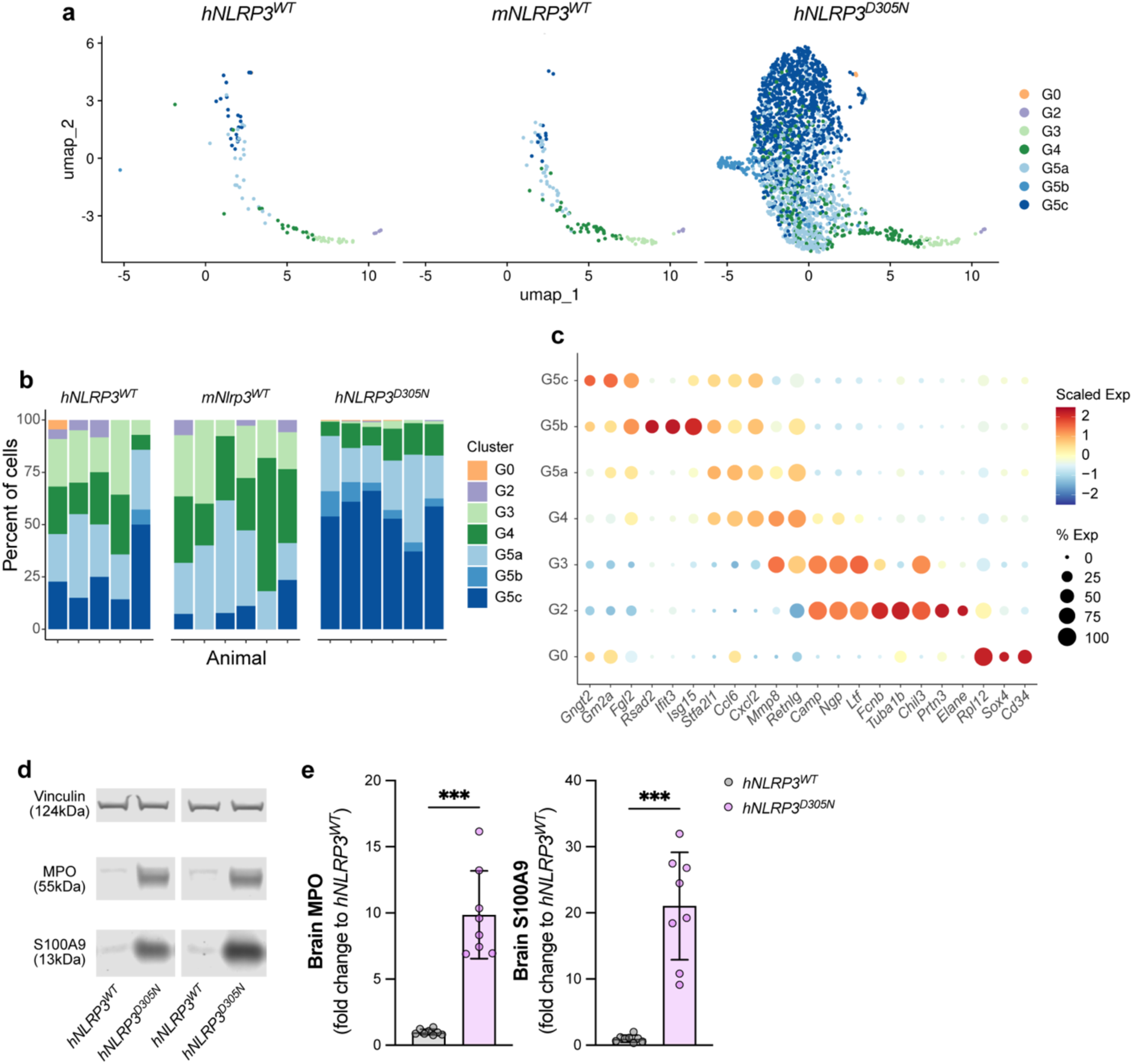
*hNLRP3^D305N^*brain neutrophils show transcriptional profiles and express proteins associated with reactivity (a-c) Neutrophil scRNA-seq data from CD45^+^ cells collected via FACS from *hNLRP3^WT^*, *mNlrp3^WT^*, and *hNLRP3^D305N^* mouse brains (Supplementary Fig. 3). Neutrophil subpopulations were assigned using the neutrophil transcriptome profiling in Xie *et al.*^62^ (n = 6/group). **(a)** UMAP plots separated by genotype, showing neutrophil subpopulations after reference mapping. **(b)** Percent of each neutrophil subpopulation per animal, separated by genotype. **(c)** scRNA-seq expression of neutrophil subpopulation marker genes from Xie *et al.*^62^ across neutrophil subpopulations. **(d-e)** Representative images and quantification of western blot for MPO and S100A9 in bulk mouse brains for *hNLRP3^WT^* and *hNLRP3^D305N^*. Vinculin used as loading control. Lognormal Welch two-tailed t-test (n = 8/group). *p < 0.05, **p < 0.01, ***p < 0.001. Data are shown as mean ± SD. Full source data and statistic values in Source Data.

**Supplementary Figure 7.**
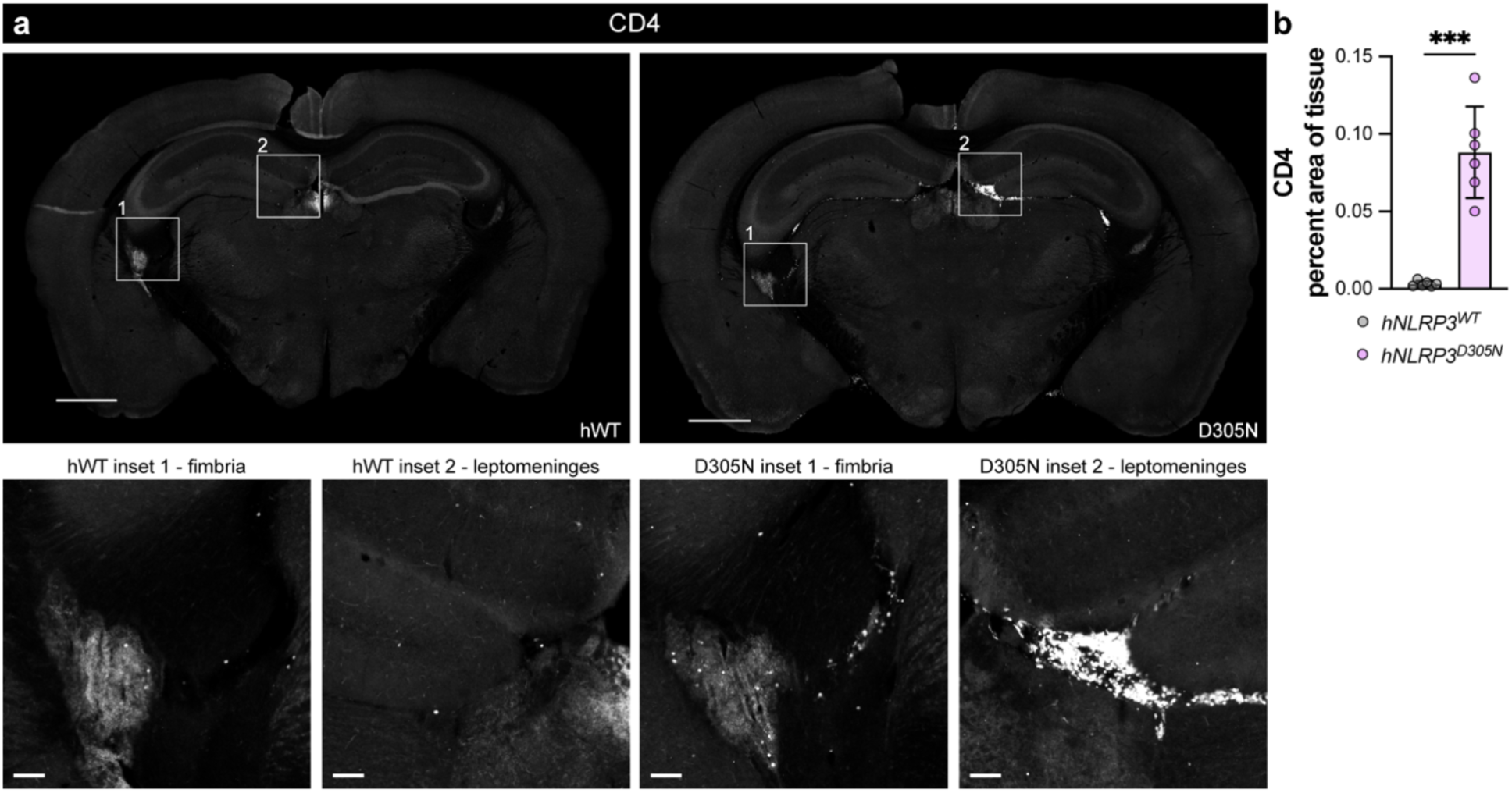
CD4^+^ T cells are elevated in *hNLRP3^D305N^* leptomeninges (a-b) Representative widefield images and quantification of CD4 IHC in mouse coronal brain sections for *hNLRP3^WT^* and *hNLRP3^D305N^*. Scale bar: whole section, 1000μm; inset, 50μm. Lognormal Welch two-tailed t-test (n = 6/group). *p < 0.05, **p < 0.01, ***p < 0.001. Data are shown as mean ± SD. Full source data and statistic values in Source Data.

**Supplementary Figure 8.**
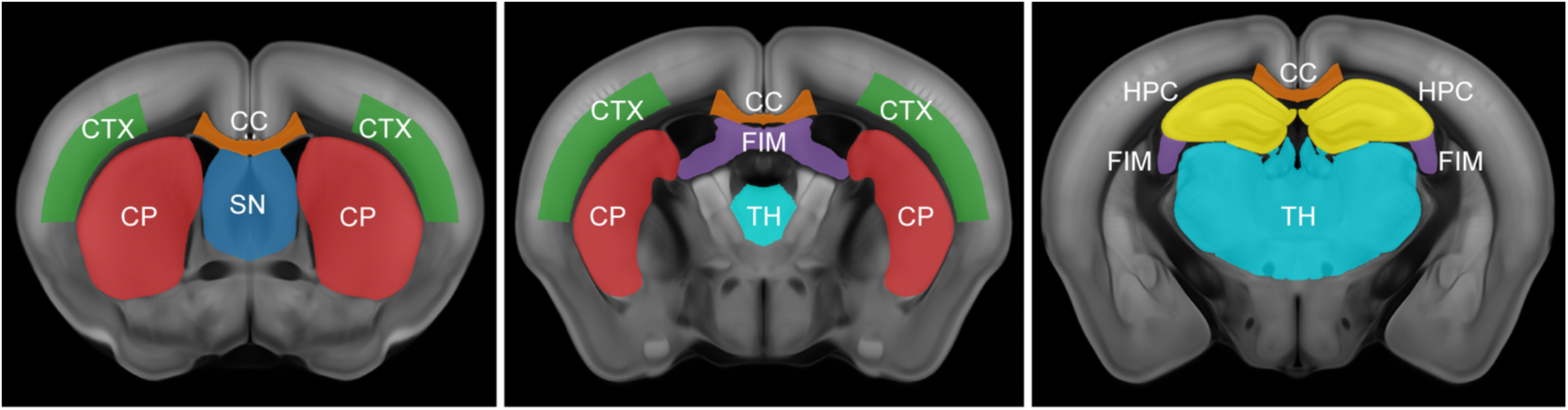
Anatomy overview of regions of interest. Anatomical regions of interest are labeled on average template coronal mouse brain sections using Allen Mouse Brain Common Coordinate Framework v3^127,128^, displayed most anterior on left to most posterior on right. CTX (green): cortex (layers 5/6 of the primary somatosensory area); CP (red): caudoputamen; CC (orange): corpus callosum (medial corpus callosum between apex of cingulum bundles); SN (blue): septal nucleus; FIM (purple): fimbria; TH (teal): thalamus; HPC (yellow): hippocampus.

**Supplementary Figure 9.**
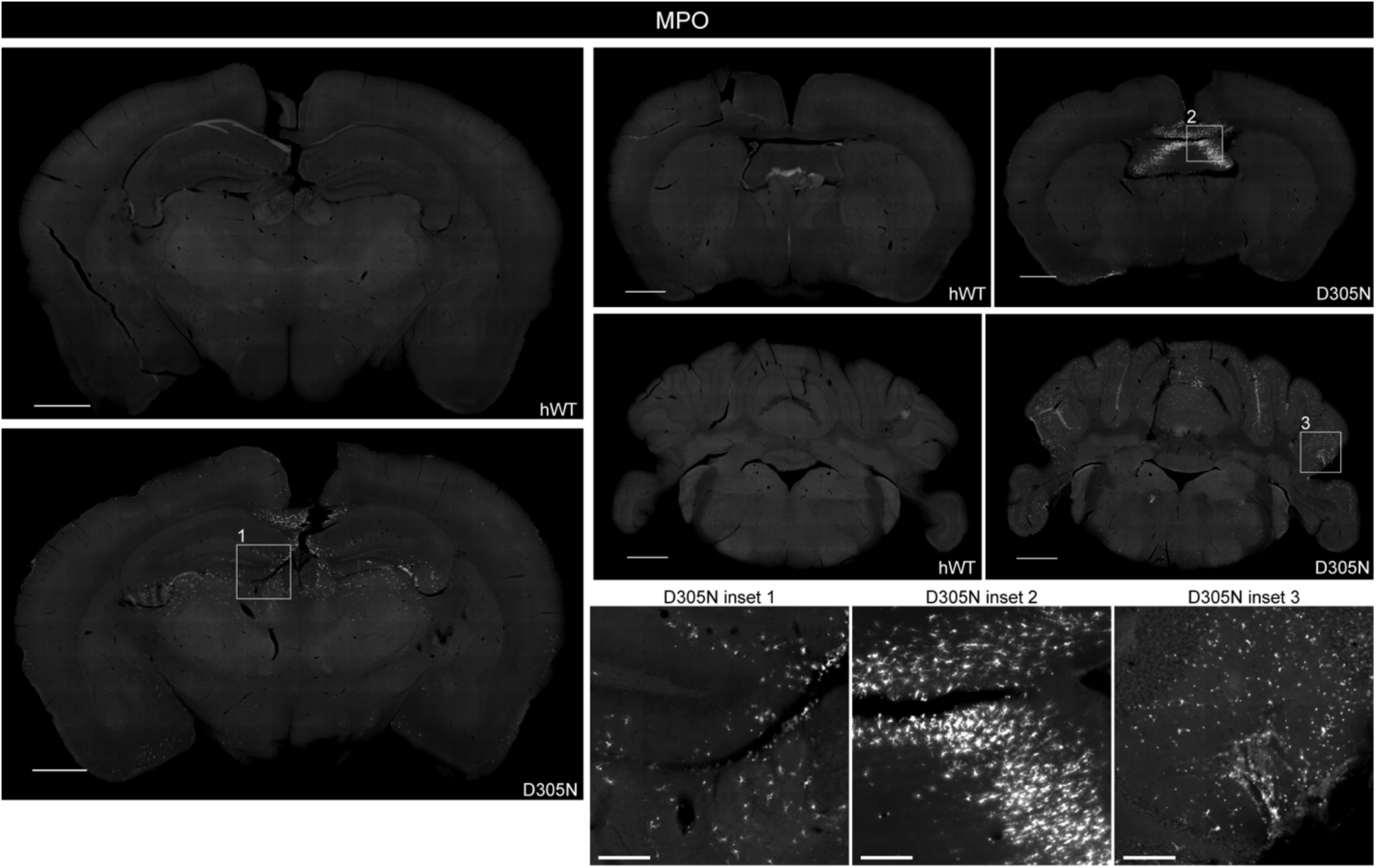
Neutrophils enter the *hNLRP3^D305N^* brain parenchyma. Representative widefield images of MPO IHC in mouse coronal brain sections. Scale bar: whole section, 1000μm; inset, 200μm.

**Supplementary Figure 10.**
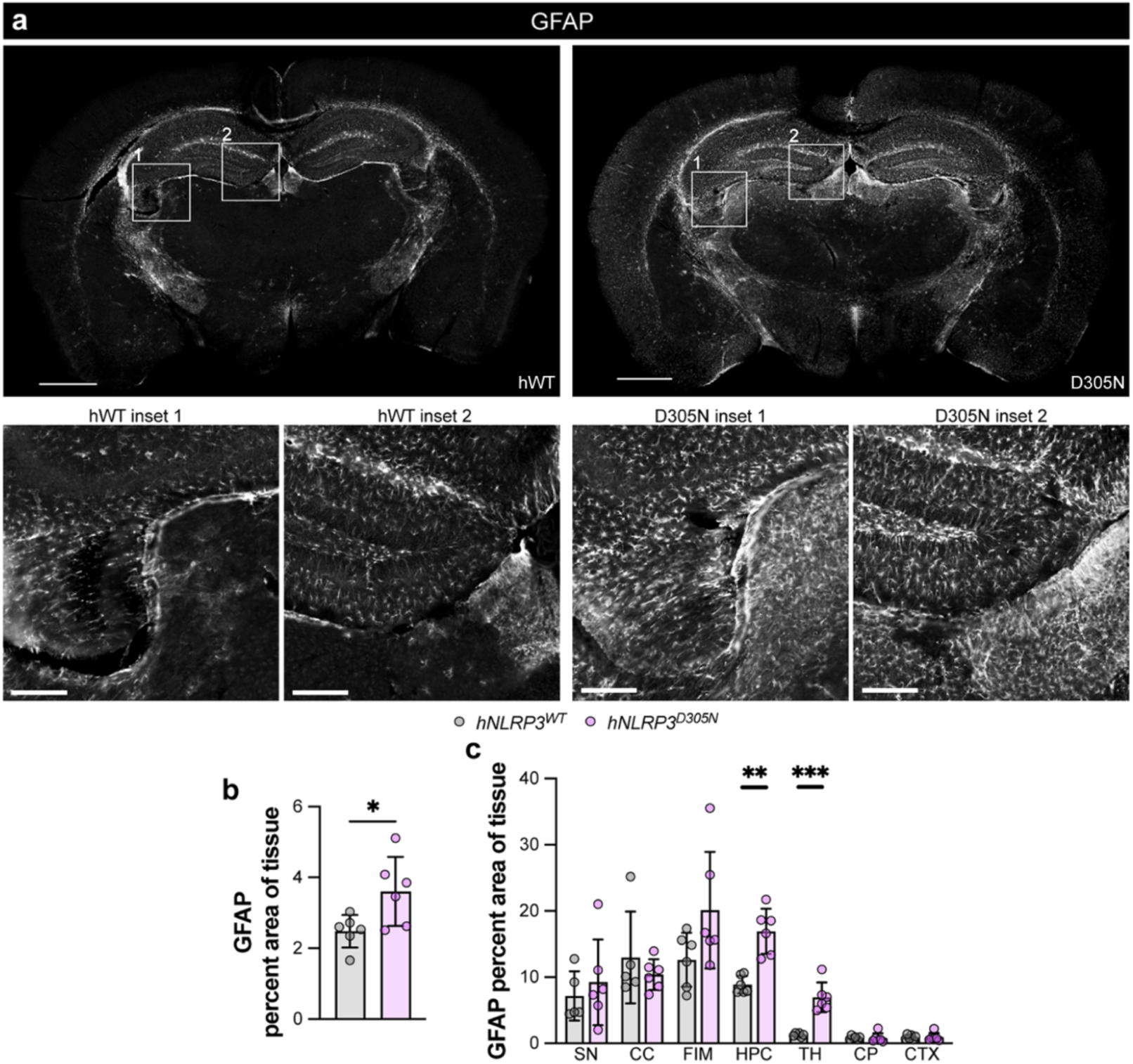
*hNLRP3^D305N^* brains show astrogliosis. *hNLRP3^WT^* and *hNLRP3^D305N^* whole brain coronal sections were stained for GFAP IHC (n = 6/group). **(a)** Representative widefield images of whole brain coronal sections. Scale bar: whole section, 1000μm; inset, 200μm. **(b)** Quantification of percent of whole tissue stained by GFAP. Lognormal Welch two-tailed t-test. **(c)** Quantification of percent of region area stained by GFAP. SN, septal nucleus; CC, corpus callosum; FIM, fimbria; HPC, hippocampus; TH, thalamus; CP, caudoputamen; CTX, cortex. Lognormal Welch two-tailed t-tests with Holm-Šídák multiple testing correction for genotype comparison in each region. *p_adj_ < 0.05, **p_adj_ < 0.01, ***p_adj_ < 0.001. Data are shown as mean ± SD. Full source data and statistic values in Source Data.

Supplementary Data 1 | Microglia pseudobulk differential gene expression results comparing *hNLRP3^D305N^*to *hNLRP3^WT^*. logFC = estimate of the log2-fold-change.

AveExpr = average log2-expression. t = moderated t-statistic. P.Value = raw p-value. adj.P.Value = adjusted p-value or q-value. B = log-odds that the gene is differentially expressed.

**Supplementary Data 2 | GSEA results for comparison of *hNLRP3^D305N^*to *hNLRP3^WT^***. pathway = name of the pathway. pval = an enrichment p-value. padj = a BH adjusted p-value. log2err = the expected error for the standard deviation of the P-value logarithm. ES = enrichment score. NES = enrichment score normalized to mean enrichment of random samples of the same size. size = size of pathway after removing genes for which contrast statistics not present.

**Supplementary Data 3 | Differential abundance statistics for brain and CSF proteomics data comparing *hNLRP3^D305N^* to *hNLRP3^WT^*.** logFC = estimate of the log2-fold-change. AveExpr = average log2-expression. t = moderated t-statistic. P.Value = raw p-value. adj.P.Value = adjusted p-value or q-value.

## References

1. Holbrook, J. A. et al. Neurodegenerative Disease and the NLRP3 Inflammasome. Front. Pharmacol. 12, 643254 (2021).

2. Anderson, F. L., Biggs, K. E., Rankin, B. E. & Havrda, M. C. NLRP3 inflammasome in neurodegenerative disease. Translational Research 252, 21–33 (2023).

3. Ravichandran, K. A. & Heneka, M. T. Inflammasomes in neurological disorders — mechanisms and therapeutic potential. Nat Rev Neurol 20, 67–83 (2024).

4. Vande Walle, L. & Lamkanfi, M. Drugging the NLRP3 inflammasome: from signalling mechanisms to therapeutic targets. Nat Rev Drug Discov 23, 43–66 (2024).

5. Cabral, J. E. et al. Targeting the NLRP3 inflammasome for inflammatory disease therapy. Trends in Pharmacological Sciences 46, 503–519 (2025).

6. Wang, H. et al. NLRP3 inflammasome in health and disease (Review). Int J Mol Med 55, 48 (2025).

7. Yao, J., Wang, Z., Song, W. & Zhang, Y. Targeting NLRP3 inflammasome for neurodegenerative disorders. Mol Psychiatry 28, 4512–4527 (2023).

8. Huang, Y., Xu, W. & Zhou, R. NLRP3 inflammasome activation and cell death. Cell Mol Immunol 18, 2114–2127 (2021).

9. Seok, J. K., Kang, H. C., Cho, Y.-Y., Lee, H. S. & Lee, J. Y. Regulation of the NLRP3 Inflammasome by Post-Translational Modifications and Small Molecules. Front. Immunol. 11, 618231 (2021).

10. Heneka, M. T., McManus, R. M. & Latz, E. Inflammasome signalling in brain function and neurodegenerative disease. Nat Rev Neurosci 19, 610–621 (2018).

11. Swanson, K. V., Deng, M. & Ting, J. P.-Y. The NLRP3 inflammasome: molecular activation and regulation to therapeutics. Nat Rev Immunol 19, 477–489 (2019).

12. Hoss, F., Rodriguez-Alcazar, J. F. & Latz, E. Assembly and regulation of ASC specks. Cell. Mol. Life Sci. 74, 1211–1229 (2017).

13. Shin, H. J., Kim, I. S., Kim, J. K. & Jo, E.-K. Molecular mechanisms of NLRP3 inflammasome activation. Exp Mol Med 58, 650–663 (2026).

14. Newton, K., Dixit, V. M. & Kayagaki, N. Dying cells fan the flames of inflammation. Science 374, 1076–1080 (2021).

15. Accogli, T., Hibos, C. & Vegran, F. Canonical and non-canonical functions of NLRP3. Journal of Advanced Research S2090123223000012 (2023) doi:10.1016/j.jare.2023.01.001.

16. Lewcock, J. W. et al. Emerging Microglia Biology Defines Novel Therapeutic Approaches for Alzheimer’s Disease. Neuron 108, 801–821 (2020).

17. Li, T., Zheng, G., Li, B. & Tang, L. Pyroptosis: A promising therapeutic target for noninfectious diseases. Cell Proliferation 54, e13137 (2021).

18. Mustafa, M. A. et al. Exploring the Role of NLRP3 in Neurodegeneration: Cutting-Edge Therapeutic Strategies and Inhibitors. Developmental Neurobiology 85, e22982 (2025).

19. Mendiola, A. S. & Cardona, A. E. The IL-1β phenomena in neuroinflammatory diseases. J Neural Transm 125, 781–795 (2018).

20. Moonen, S. et al. Pyroptosis in Alzheimer’s disease: cell type-specific activation in microglia, astrocytes and neurons. Acta Neuropathol 145, 175–195 (2023).

21. Kadhim, H., Deltenre, P., Martin, J.-J. & Sébire, G. In-situ expression of Interleukin-18 and associated mediators in the human brain of sALS patients: Hypothesis for a role for immune-inflammatory mechanisms. Medical Hypotheses 86, 14–17 (2016).

22. Gordon, R. et al. Inflammasome inhibition prevents α-synuclein pathology and dopaminergic neurodegeneration in mice. Science Translational Medicine 10, eaah4066 (2018).

23. Piancone, F., La Rosa, F., Marventano, I., Saresella, M. & Clerici, M. The Role of the Inflammasome in Neurodegenerative Diseases. Molecules 26, 953 (2021).

24. Heneka, M. T. et al. NLRP3 is activated in Alzheimer’s disease and contributes to pathology in APP/PS1 mice. Nature 493, 674–678 (2013).

25. Venegas, C. et al. Microglia-derived ASC specks cross-seed amyloid-β in Alzheimer’s disease. Nature 552, 355–361 (2017).

26. Ising, C. et al. NLRP3 inflammasome activation drives tau pathology. Nature 575, 669–673 (2019).

27. Lonnemann, N. et al. The NLRP3 inflammasome inhibitor OLT1177 rescues cognitive impairment in a mouse model of Alzheimer’s disease. Proc. Natl. Acad. Sci. U.S.A. 117, 32145–32154 (2020).

28. Qiao, C. et al. Inhibition of the hepatic Nlrp3 protects dopaminergic neurons via attenuating systemic inflammation in a MPTP/p mouse model of Parkinson’s disease. J Neuroinflammation 15, 193 (2018).

29. Huang, S. et al. A selective NLRP3 inflammasome inhibitor attenuates behavioral deficits and neuroinflammation in a mouse model of Parkinson’s disease. Journal of Neuroimmunology 354, 577543 (2021).

30. Panicker, N. et al. Neuronal NLRP3 is a parkin substrate that drives neurodegeneration in Parkinson’s disease. Neuron 110, 2422–2437.e9 (2022).

31. Deora, V. et al. The microglial NLRP3 inflammasome is activated by amyotrophic lateral sclerosis proteins. Glia 68, 407–421 (2020).

32. Shu, X. et al. Negative regulation of TREM2-mediated C9orf72 poly-GA clearance by the NLRP3 inflammasome. Cell Reports 42, 112133 (2023).

33. Cihankaya, H. et al. Elevated NLRP3 Inflammasome Activation Is Associated with Motor Neuron Degeneration in ALS. Cells 13, 995 (2024).

34. Gris, D. et al. NLRP3 Plays a Critical Role in the Development of Experimental Autoimmune Encephalomyelitis by Mediating Th1 and Th17 Responses. The Journal of Immunology 185, 974–981 (2010).

35. Inoue, M., Williams, K. L., Gunn, M. D. & Shinohara, M. L. NLRP3 inflammasome induces chemotactic immune cell migration to the CNS in experimental autoimmune encephalomyelitis. Proc. Natl. Acad. Sci. U.S.A. 109, 10480–10485 (2012).

36. Coll, R. C. et al. A small-molecule inhibitor of the NLRP3 inflammasome for the treatment of inflammatory diseases. Nat Med 21, 248–255 (2015).

37. Hou, B. et al. Inhibiting the NLRP3 Inflammasome with MCC950 Alleviates Neurological Impairment in the Brain of EAE Mice. Mol Neurobiol 61, 1318–1330 (2024).

38. Booshehri, L. M. & Hoffman, H. M. CAPS and NLRP3. J Clin Immunol 39, 277–286 (2019).

39. Mamoudjy, N., Maurey, H., Marie, I., Koné-Paut, I. & Deiva, K. Neurological outcome of patients with cryopyrin-associated periodic syndrome (CAPS). Orphanet J Rare Dis 12, 33 (2017).

40. Lee, J., Cho, W., Yu, J.-W. & Hyun, Y.-M. NLRP3 activation induces BBB disruption and neutrophil infiltration via CXCR2 signaling in the mouse brain. J Neuroinflammation 22, 139 (2025).

41. Snouwaert, J. N. et al. An NLRP3 Mutation Causes Arthropathy and Osteoporosis in Humanized Mice. Cell Reports 17, 3077–3088 (2016).

42. Koller, B. H., Nguyen, M., Snouwaert, J. N., Gabel, C. A. & Ting, J. P.-Y. Species-specific NLRP3 regulation and its role in CNS autoinflammatory diseases. Cell Reports 43, (2024).

43. Koller, B. H. et al. CNS-targeted NLRP3 Inhibition by NT-0527 confers therapeutic advantage in a CAPS mouse model. J Neuroinflammation 23, 104 (2026).

44. Snouwaert, J. N., et al. Mice humanized by syntenic replacement with full-length NLRP3 disease-associated variants model the clinical cryopyrinopathy continuum. JCI Insight 11, e194677 (2026).

45. Arend, W. P. The balance between IL-1 and IL-1Ra in disease. Cytokine & Growth Factor Reviews 13, 323–340 (2002).

46. Yoon, S.-H. et al. Microglial NLRP3-gasdermin D activation impairs blood-brain barrier integrity through interleukin-1β-independent neutrophil chemotaxis upon peripheral inflammation in mice. Nat Commun 16, 699 (2025).

47. Yang, J. et al. New insight into neurological degeneration: Inflammatory cytokines and blood–brain barrier. Front. Mol. Neurosci. 15, 1013933 (2022).

48. Logan, T. et al. Rescue of a lysosomal storage disorder caused by Grn loss of function with a brain penetrant progranulin biologic. Cell 184, 4651–4668.e25 (2021).

49. Muenzer, J. et al. Cerebrospinal fluid heparan sulfate as a biomarker for neuronopathic mucopolysaccharidoses: Rationale and regulatory challenges. Molecular Genetics and Metabolism 148, 109911 (2026).

50. Siletti, K. et al. Transcriptomic diversity of cell types across the adult human brain. Science 382, eadd7046 (2023).

51. Single cell type - NLRP3 - The Human Protein Atlas. https://www.proteinatlas.org/ENSG00000162711-NLRP3/single+cell#immune_cell.

52. Butovsky, O. & Weiner, H. L. Microglial signatures and their role in health and disease. Nat Rev Neurosci 19, 622–635 (2018).

53. Keren-Shaul, H. et al. A Unique Microglia Type Associated with Restricting Development of Alzheimer’s Disease. Cell 169, 1276–1290.e17 (2017).

54. Fumagalli, L. et al. Microglia heterogeneity, modeling and cell-state annotation in development and neurodegeneration. Nat Neurosci 28, 1381–1392 (2025).

55. Rosmus, D.-D., Lange, C., Ludwig, F., Ajami, B. & Wieghofer, P. The Role of Osteopontin in Microglia Biology: Current Concepts and Future Perspectives. Biomedicines 10, 840 (2022).

56. Bhattarai, P. et al. Rare genetic variation in fibronectin 1 (FN1) protects against APOEε4 in Alzheimer’s disease. Acta Neuropathol 147, 70 (2024).

57. Chen, K., Tang, W. & Liu, X. Research and progress of cGAS/STING/NLRP3 signaling pathway: a mini review. Front. Immunol. 16, 1594133 (2025).

58. Krasemann, S. et al. The TREM2-APOE Pathway Drives the Transcriptional Phenotype of Dysfunctional Microglia in Neurodegenerative Diseases. Immunity 47, 566–581.e9 (2017).

59. Paolicelli, R. C. et al. Microglia states and nomenclature: A field at its crossroads. Neuron 110, 3458–3483 (2022).

60. Karlsson, M. et al. A single–cell type transcriptomics map of human tissues. SCIENCE ADVANCES (2021).

61. Paget, C., Doz-Deblauwe, E., Winter, N. & Briard, B. Specific NLRP3 Inflammasome Assembling and Regulation in Neutrophils: Relevance in Inflammatory and Infectious Diseases. Cells 11, 1188 (2022).

62. Xie, X. et al. Single-cell transcriptome profiling reveals neutrophil heterogeneity in homeostasis and infection. Nat Immunol 21, 1119–1133 (2020).

63. Urban, C. F. et al. Neutrophil Extracellular Traps Contain Calprotectin, a Cytosolic Protein Complex Involved in Host Defense against Candida albicans. PLoS Pathog 5, e1000639 (2009).

64. Zhou, Y. et al. Role of S100A8/A9 for Cytokine Secretion, Revealed in Neutrophils Derived from ER-Hoxb8 Progenitors. IJMS 22, 8845 (2021).

65. Sprenkeler, E. G. G. et al. S100A8/A9 Is a Marker for the Release of Neutrophil Extracellular Traps and Induces Neutrophil Activation. Cells 11, 236 (2022).

66. Romejko, K., Markowska, M. & Niemczyk, S. The Review of Current Knowledge on Neutrophil Gelatinase-Associated Lipocalin (NGAL). IJMS 24, 10470 (2023).

67. Liu, Y. et al. Lcn2 from neutrophil extracellular traps induces astrogliosis and post-stroke emotional disorders. Neuron 113, 4199–4216.e8 (2025).

68. Shafqat, A. et al. Neutrophil extracellular traps in central nervous system pathologies: A mini review. Front. Med. 10, 1083242 (2023).

69. Wang, H. et al. Neutrophil extracellular traps in homeostasis and disease. Sig Transduct Target Ther 9, 235 (2024).

70. Leinweber, J. et al. Elastase inhibitor agaphelin protects from acute ischemic stroke in mice by reducing thrombosis, blood–brain barrier damage, and inflammation. Brain, Behavior, and Immunity 93, 288–298 (2021).

71. Villalba, N., Baby, S., Cha, B. J. & Yuan, S. Y. Site-specific opening of the blood-brain barrier by extracellular histones. J Neuroinflammation 17, 281 (2020).

72. Silvestre-Roig, C. et al. Externalized histone H4 orchestrates chronic inflammation by inducing lytic cell death. Nature 569, 236–240 (2019).

73. Saffarzadeh, M. et al. Neutrophil Extracellular Traps Directly Induce Epithelial and Endothelial Cell Death: A Predominant Role of Histones. PLoS ONE 7, e32366 (2012).

74. Santos-Lima, B., Pietronigro, E. C., Terrabuio, E., Zenaro, E. & Constantin, G. The role of neutrophils in the dysfunction of central nervous system barriers. Front. Aging Neurosci. 14, 965169 (2022).

75. Strzepa, A., Pritchard, K. A. & Dittel, B. N. Myeloperoxidase: A new player in autoimmunity. Cellular Immunology 317, 1–8 (2017).

76. Sankowski, R. et al. Multiomic spatial landscape of innate immune cells at human central nervous system borders. Nat Med (2023) doi:10.1038/s41591-023-02673-1.

77. Betsholtz, C. et al. Advances and controversies in meningeal biology. Nat Neurosci 27, 2056–2072 (2024).

78. Parker, T. et al. Neurology of the cryopyrin-associated periodic fever syndrome. Euro J of Neurology 23, 1145–1151 (2016).

79. Fleming, T. J., Fleming, M. L. & Malek, T. R. Selective expression of Ly-6G on myeloid lineage cells in mouse bone marrow. RB6-8C5 mAb to granulocyte-differentiation antigen (Gr-1) detects members of the Ly-6 family. J Immunol 151, 2399–2408 (1993).

80. Mercurio, D. et al. Protein Expression of the Microglial Marker Tmem119 Decreases in Association With Morphological Changes and Location in a Mouse Model of Traumatic Brain Injury. Front. Cell. Neurosci. 16, 820127 (2022).

81. Leyh, J. et al. Classification of Microglial Morphological Phenotypes Using Machine Learning. Front. Cell. Neurosci. 15, 701673 (2021).

82. Green, T. R. F. & Rowe, R. K. Quantifying microglial morphology: an insight into function. Clinical and Experimental Immunology 216, 221–229 (2024).

83. Green, T. R. F., Murphy, S. M. & Rowe, R. K. Comparisons of quantitative approaches for assessing microglial morphology reveal inconsistencies, ecological fallacy, and a need for standardization. Sci Rep 12, 18196 (2022).

84. Reymann, J. LIGHTNING: Image Information Extraction By Adaptive Deconvolution. (2018).

85. Giovannoni, F. & Quintana, F. J. The Role of Astrocytes in CNS Inflammation. Trends in Immunology 41, 805–819 (2020).

86. Matejuk, A. & Ransohoff, R. M. Crosstalk Between Astrocytes and Microglia: An Overview. Front. Immunol. 11, 1416 (2020).

87. Escartin, C. et al. Reactive astrocyte nomenclature, definitions, and future directions. Nat Neurosci 24, 312–325 (2021).

88. Ardic, N. & Dinc, R. Neutrophil extracellular traps and microglia/macrophages interactions in stroke: from thromboinflammation to immunotherapy. Front. Immunol. 17, 1752471 (2026).

89. Neumann, J. et al. Microglia Cells Protect Neurons by Direct Engulfment of Invading Neutrophil Granulocytes: A New Mechanism of CNS Immune Privilege. Journal of Neuroscience 28, 5965–5975 (2008).

90. Neumann, J. et al. Beware the intruder: Real time observation of infiltrated neutrophils and neutrophil—Microglia interaction during stroke in vivo. PLoS ONE 13, e0193970 (2018).

91. Sollberger, G., Tilley, D. O. & Zychlinsky, A. Neutrophil Extracellular Traps: The Biology of Chromatin Externalization. Developmental Cell 44, 542–553 (2018).

92. Palma, A. The Landscape of SPP1^+^ Macrophages Across Tissues and Diseases: A Comprehensive Review. Immunology 176, 179–196 (2025).

93. Wang, H. et al. FN1 is a prognostic biomarker and correlated with immune infiltrates in gastric cancers. Front. Oncol. 12, 918719 (2022).

94. Hoeft, K. et al. Platelet-instructed SPP1+ macrophages drive myofibroblast activation in fibrosis in a CXCL4-dependent manner. Cell Reports 42, 112131 (2023).

95. Monroe, K. M., Hong, S., Lewcock, J. W. & Yang, A. C. Therapeutic targeting of neuroimmune mechanisms in neurodegeneration. Nat Rev Drug Discov (2026) doi:10.1038/s41573-025-01370-7.

96. Sica, A., Matsushima, K. & Colotta, F. IL-1 transcriptionally activates the neutrophil chemotactic factor/IL-8 gene in endothelial cells. (1990).

97. Ebisawa, M., Bochner, B. S., Georas, S. N. & Schleimer, R. P. Eosinophil transendothelial migration induced by cytokines. I. Role of endothelial and eosinophil adhesion molecules in IL-1 beta-induced transendothelial migration. The Journal of Immunology 149, 4021–4028 (1992).

98. Shaftel, S. S. et al. Chronic Interleukin-1β Expression in Mouse Brain Leads to Leukocyte Infiltration and Neutrophil-Independent Blood–Brain Barrier Permeability without Overt Neurodegeneration. J. Neurosci. 27, 9301–9309 (2007).

99. Labus, J. et al. IL-1β promotes transendothelial migration of PBMCs by upregulation of the FN/α5β1 signalling pathway in immortalised human brain microvascular endothelial cells. Experimental Cell Research 373, 99–111 (2018).

100. Eyles, J. L. et al. A key role for G-CSF–induced neutrophil production and trafficking during inflammatory arthritis. Blood 112, 5193–5201 (2008).

101. Summers, C. et al. Neutrophil kinetics in health and disease. Trends in Immunology 31, 318–324 (2010).

102. Rossi, B., Constantin, G. & Zenaro, E. The emerging role of neutrophils in neurodegeneration. Immunobiology 225, 151865 (2020).

103. Wright, H. L., Cross, A. L., Edwards, S. W. & Moots, R. J. Effects of IL-6 and IL-6 blockade on neutrophil function in vitro and in vivo. Rheumatology 53, 1321–1331 (2014).

104. Florentin, J. et al. Interleukin-6 mediates neutrophil mobilization from bone marrow in pulmonary hypertension. Cell Mol Immunol 18, 374–384 (2021).

105. Zhang, F. et al. Neutrophil diversity and function in health and disease. Sig Transduct Target Ther 9, 343 (2024).

106. Capucetti, A., Albano, F. & Bonecchi, R. Multiple Roles for Chemokines in Neutrophil Biology. Front. Immunol. 11, 1259 (2020).

107. Sawant, K. V. et al. Neutrophil recruitment by chemokines Cxcl1/KC and Cxcl2/MIP2: Role of Cxcr2 activation and glycosaminoglycan interactions. Journal of Leukocyte Biology 109, 777–791 (2021).

108. Onishi, R. M. & Gaffen, S. L. Interleukin-17 and its target genes: mechanisms of interleukin-17 function in disease. Immunology 129, 311–321 (2010).

109. Fan, X., Shu, P., Wang, Y., Ji, N. & Zhang, D. Interactions between neutrophils and T-helper 17 cells. Front. Immunol. 14, 1279837 (2023).

110. Rosenzweig, N. et al. Sex-dependent APOE4 neutrophil–microglia interactions drive cognitive impairment in Alzheimer’s disease. Nat Med (2024) doi:10.1038/s41591-024-03122-3.

111. Mei, J. et al. Cxcr2 and Cxcl5 regulate the IL-17/G-CSF axis and neutrophil homeostasis in mice. J. Clin. Invest. 122, 974–986 (2012).

112. Wang, L.-Y., Tu, Y.-F., Lin, Y.-C. & Huang, C.-C. CXCL5 signaling is a shared pathway of neuroinflammation and blood–brain barrier injury contributing to white matter injury in the immature brain. J Neuroinflammation 13, 6 (2016).

113. Kolaczkowska, E. & Kubes, P. Neutrophil recruitment and function in health and inflammation. Nat Rev Immunol 13, 159–175 (2013).

114. Bonar, S. L. et al. Constitutively Activated NLRP3 Inflammasome Causes Inflammation and Abnormal Skeletal Development in Mice. PLoS ONE 7, e35979 (2012).

115. Schonhoff, A. M. et al. Border-associated macrophages mediate the neuroinflammatory response in an alpha-synuclein model of Parkinson disease. Nat Commun 14, 3754 (2023).

116. Hobson, R. et al. Clonal CD8+ T cells populate the leptomeninges and coordinate with immune cells in human degenerative brain diseases. Nat Immunol 27, 323–335 (2026).

117. Xiong, J., Xue, J., Zhou, H., Qi, W. & Zhu, H. The crosstalk of neutrophil extracellular trap-inflammasome and their roles in disease progression. Molecular Aspects of Medicine 106, 101419 (2025).

118. Wang, L. et al. Targeting the interactions between neutrophils and microglia: a novel strategy for anti-inflammatory treatment of stroke. Acta Pharmacol Sin 47, 549–565 (2026).

119. Liu, L. et al. NLRP3/GSDMD-dependent neutrophil extracellular traps exacerbate microglia-mediated neuroinflammation following traumatic brain injury. Cell Commun Signal 24, 65 (2026).

120. Qin, L. et al. Systemic LPS causes chronic neuroinflammation and progressive neurodegeneration. Glia 55, 453–462 (2007).

121. Bai, B. et al. NLRP3 inflammasome in endothelial dysfunction. Cell Death Dis 11, 776 (2020).

122. Johann, S. et al. NLRP3 inflammasome is expressed by astrocytes in the SOD1 mouse model of ALS and in human sporadic ALS patients: NLRP3 Inflammasome Expression in ALS. Glia 63, 2260–2273 (2015).

123. Ullman, J. C. et al. Brain delivery and activity of a lysosomal enzyme using a blood-brain barrier transport vehicle in mice. Sci. Transl. Med. 12, eaay1163 (2020).

124. Meyer, T. et al. Neurofilament light-chain response during therapy with antisense oligonucleotide tofersen in SOD1-related ALS: Treatment experience in clinical practice. Muscle and Nerve 67, 515–521 (2023).

125. Smith, S. E. et al. Tofersen treatment leads to sustained stabilization of disease in SOD1 ALS in a “real-world” setting. Ann Clin Transl Neurol 12, 311–319 (2025).

126. Gonzalez, G. et al. CSF turnover reshapes biomarker interpretation in neurodegeneration studies. Preprint at 10.64898/2026.02.02.26345363 (2026).

127. Reference Atlas:: Allen Brain Atlas: Mouse Brain. https://mouse.brain-map.org/static/atlas.

128. Wang, Q. et al. The Allen Mouse Brain Common Coordinate Framework: A 3D Reference Atlas. Cell 181, 936–953.e20 (2020).

129. van Lengerich, B. et al. A TREM2-activating antibody with a blood–brain barrier transport vehicle enhances microglial metabolism in Alzheimer’s disease models. Nat Neurosci (2023) doi:10.1038/s41593-022-01240-0.

130. Van Der Walt, S. et al. scikit-image: image processing in Python. PeerJ 2, e453 (2014).

131. Virtanen, P. et al. SciPy 1.0: fundamental algorithms for scientific computing in Python. Nat Methods 17, 261–272 (2020).

132. Pedregosa, F. et al. Scikit-learn: Machine Learning in Python. MACHINE LEARNING IN PYTHON (2011).

133. R: The R Project for Statistical Computing. https://www.r-project.org/.

134. Hao, Y. et al. Integrated analysis of multimodal single-cell data. Cell 184, 3573–3587.e29 (2021).

135. Hao, Y. et al. Dictionary learning for integrative, multimodal and scalable single-cell analysis. Nat Biotechnol 42, 293–304 (2024).

136. Germain, P.-L., Lun, A., Meixide, C. G., Macnair, W. & Robinson, M. D. Doublet identification in single-cell sequencing data using scDblFinder. (2022) 10.12688/f1000research.73600.1.

137. Wickham, H. Ggplot2: Elegant Graphics for Data Analysis. (Springer-Verlag New York, 2016).

138. de Vries, A. & Ripley, B. D. ggdendro: Create Dendrograms and Tree Diagrams Using ‘ggplot2’. R package version 0.2.0. (2025).

139. Wilke, C. O. cowplot: Streamlined Plot Theme and Plot Annotations for ‘ggplot2’. R package version 1.2.0. (2025).

140. Zappia, L. & Oshlack, A. Clustering trees: a visualization for evaluating clusterings at multiple resolutions. Gigascience 7, giy083 (2018).

141. McCarthy, D. J., Campbell, K. R., Lun, A. T. L. & Wills, Q. F. Scater: pre-processing, quality control, normalization and visualization of single-cell RNA-seq data in R. Bioinformatics 33, 1179–1186 (2017).

142. Ritchie, M. E. et al. limma powers differential expression analyses for RNA-sequencing and microarray studies. Nucleic Acids Res 43, e47 (2015).

143. Chen, Y., Chen, L., Lun, A. T. L., Baldoni, P. L. & Smyth, G. K. edgeR v4: powerful differential analysis of sequencing data with expanded functionality and improved support for small counts and larger datasets. Nucleic Acids Res 53, gkaf018 (2025).

144. Subramanian, A. et al. Gene set enrichment analysis: a knowledge-based approach for interpreting genome-wide expression profiles. Proc Natl Acad Sci U S A 102, 15545–15550 (2005).

145. Korotkevich, G. et al. Fast gene set enrichment analysis. 060012 Preprint at 10.1101/060012 (2021).

146. Liberzon, A. et al. The Molecular Signatures Database (MSigDB) hallmark gene set collection. Cell Syst 1, 417–425 (2015).

147. Dolgalev, I. msigdbr: MSigDB Gene Sets for Multiple Organisms in a Tidy Data Format. R package version 26.1.0. (2026).

148. Korsunsky, I. et al. Fast, sensitive and accurate integration of single-cell data with Harmony. Nat Methods 16, 1289–1296 (2019).

149. Korsunsky, I. et al. _harmony: Fast, Sensitive, and Accurate Integration of Single Cell Data_. R package version 1.2.4. (2026).

150. Stuart, T. et al. Comprehensive Integration of Single-Cell Data. Cell 177, 1888–1902.e21 (2019).

151. Yulyaningsih, E. et al. DNL343 is an investigational CNS penetrant eukaryotic initiation factor 2B activator that prevents and reverses the effects of neurodegeneration caused by the integrated stress response. eLife 12, RP92173 (2024).

152. Davis, O. B. et al. A Common PD-Risk GBA1 Variant Disrupts LIMP2 Interaction, Impairs Glucocerebrosidase Function, and Drives Lysosomal and Mitochondrial Dysfunction. bioRxiv 2025.08.28.672891 (2025) doi:10.1101/2025.08.28.672891.

153. Motulsky, H. J., Head, T. & Clarke, P. B. S. Analyzing lognormal data: A nonmathematical practical guide. Pharmacological Reviews 77, 100049 (2025).

